# Frataxin depletion leads to decreased soma size and activation of AMPK metabolic pathway in dorsal root ganglia sensory neurons

**DOI:** 10.1101/2025.10.07.680891

**Authors:** Olivier Griso, Deepika Chellapandi, Amélie Weiss, Ioannis Manolaras, Hélène Puccio

## Abstract

**Introduction:** Friedreich’s ataxia (FA) is an inherited neurodegenerative disorder caused by frataxin deficiency, leading to impaired iron–sulfur (Fe–S) cluster biogenesis and mitochondrial dysfunction. Proprioceptive dorsal root ganglia (DRG) neurons are among the most vulnerable cell types in FA, yet the mechanisms underlying their selective susceptibility remain unclear.

**Methods:** We developed a primary culture model of embryonic mouse DRG neurons with complete frataxin depletion, enabling longitudinal analysis of cellular and metabolic alterations.

**Results:** This novel model reproduces several key biochemical hallmarks of FA, including Fe–S enzyme deficiency, mitochondrial dysfunction, altered iron homeostasis and oxidative stress. Despite severe mitochondrial impairment, frataxin-deficient neurons remained viable but exhibited a marked reduction in soma growth. Mechanistically, this phenotype was associated with activation of AMP-activated protein kinase (AMPK) hyperactivation and inhibition of mTOR signaling. Genetics inhibition of AMPK or re-expression of frataxin restored neuronal growth, demonstrating a functional link between metabolic stress signaling and soma size regulation. Treatment with α-lipoic acid (ALA) rescued soma growth, normalize ATP levels and reduce AMPK activation.

**Conclusions:** Our findings identify AMPK–mTOR signaling as a key pathway linking mitochondrial dysfunction to impaired neuronal growth impairment in frataxin-deficient sensory neurons. This robust, scalable cellular model provides new insights into sensory neuron vulnerability and offers a platform for therapeutic discovery.

## Background

Friedreich’s Ataxia (FA) is an early-onset, progressive neurodegenerative disease characterized by mixed sensory and spinocerebellar ataxia associated to hypertrophic cardiomyopathy and increased incidence of diabetes mellitus(1–3). In most case, FA is caused by a homozygous GAA trinucleotide repeat expansion within the first intron of the frataxin gene (*FXN*)(4). This expansion promotes heterochromatin formation and the generation of non-canonical DNA structures(5), leading to transcriptional repression of *FXN* and a consequent reduction in frataxin protein levels to approximately 5-30% of normal (3). FXN is a mitochondrial protein essential for the biogenesis of iron-sulfur clusters (Fe-S), key cofactors in numerous cellular processes (6). Accordingly, frataxin deficiency leads to a primary loss of Fe-S synthesis, impairs mitochondrial dysfunction, and perturbs iron metabolism, as observed in patient tissues and a variety of cellular and animal models (7–10). Increased sensitivity to oxidative stress has also been reported in several models (8,11). Thus, mitochondrial dysfunction and reactive oxygen species (ROS) accumulation are thought to be central features of the pathophysiology of FA.

The hallmark of sensory ataxia in FA primarily results from degeneration of the large, myelinated proprioceptive neurons (PNs) located in the dorsal root ganglia (DRG), key component of the sensory system responsible for transmitting information about body position and movement(12). Although traditionally regarded as a neurodegenerative disease, accumulating evidence points to a significant developmental component, particularly in the sensory system. DRG pathology emerges early in the disease course, possibly preceding symptoms onset, and shows limited progression over time. Neuropathological studies of FA patients reveal hypoplasia of both the DRG and spinal cord, and in some cases, failure of dorsal root fibers to properly enter the spinal cord (13,14). These findings suggest a failure of sensory system development. Nevertheless, postmortem analyses of patients with long disease duration have also identified signs of active neuroinflammation in the DRG, indicative of ongoing degeneration(15). Taken together, these observations support a model in which both developmental deficit and progressive neurodegeneration contribute to DRG pathology in FA. The selective vulnerability of PNs in FA remains incompletely understood, particularly given the broad tissue expression of frataxin. However, frataxin is developmentally regulated and peaks during critical stages of sensory neuron differentiation and maturation in the mouse embryo(16). Within the mature DRG, frataxin expression is highest in the PNs, correlating with their susceptibility to disease degeneration(17).

A variety of cellular models have been established to explore the effect of frataxin deficiency in sensory neurons. Primary DRG cultures from adult mice carrying a GAA expansion or from rat with frataxin knockdown recapitulate aspects of FA pathology, including mitochondrial dysfunction, impaired neurite outgrowth, altered growth-cone dynamics, and disrupted calcium signaling(18–20). More recently, iPSC-derived sensory neurons from FA patients have been shown to exhibit downregulation of transcription factors involved in PN specification and survival, abnormal neurite processes, delayed electrophysiological maturation, and transcriptomic dysregulation affecting cytoskeletal organization, neurite extension, synaptic plasticity, and neurotransmission(21–23). While these models support the notion of both developmental impairment and neurodegeneration, the molecular pathways underlying these phenotypes remain insufficiently defined. Furthermore, the phenotypes observed in these systems tend to be modest and variable, limiting their utility for mechanistic dissection and high-throughput screening (HTS).

To overcome these limitations, we developed an *in vitro* model of FA based on primary cultures of embryonic mouse DRG sensory neurons carrying a conditional *Fxn* allele(24). Upon Cre-mediated excision, these cultures exhibit complete frataxin deficiency, enabling longitudinal studies of mitochondrial function, oxidative stress, and cellular morphology. Although more severe than partial deficiency, this model recapitulates key biochemical features of FA, including Fe-S deficiency, mitochondrial iron accumulation, altered bioenergetics, and sensitivity to oxidative stress. In the present study, we use this model to investigate the impact of FXN loss on the metabolic and morphological integrity of DRG sensory neurons. We found that frataxin-deficient neurons remain viable for extended periods despite severe mitochondrial dysfunction but display a pronounced reduction in soma size. Mechanistically, we identified overactivation of the AMP-activated protein kinase (AMPK) and suppression of mTOR signaling as central mediators of this growth defect. In addition, we demonstrated that FXN deficiency compromises lipoic acid-dependent enzymatic pathways and that supplementation with alpha-lipoic acid (ALA) restores neuronal growth through both metabolic and antioxidant mechanisms. Collectively, our findings establish a mechanistic link between frataxin depletion, dysregulated metabolic signaling, and impaired neuronal growth, and provide a robust platform for therapeutic screening in FA.

## Methods

### Establishment of a mouse primary DRG sensory neurons culture, infections and treatments

#### Neuronal cultures

##### Culture media

- C-medium: Minimum Essential Medium (MEM), 10% Fetal Bovine Serum (FBS), 1% Penicillin/Streptomycin (P/S), 1% L-Glutamine, 4g/L of D-Glucose, 50 nM NGF (N-100, Alomone labs).
- NB medium: Neurobasal without Glucose and pyruvate (Gibco A2477501), 2% B-27^™^ Supplement (17504044, ThermoFisher), 1% Penicillin/Streptomycin (P/S), 1% L-Glutamine, 4g/L of D-Glucose, 50 nM NGF (N-100, Alomone labs).

Mouse primary DRG sensory neurons were cultures as recently published (24). 96-well plates were coated with poly-L-lysine (0.1 mg/ml; Merck, P5899), laminin (3.3 ug/ml; ThermoFisher Scientific 23017-015), and Matrigel (1/40 dilution; Corning BV 354277). DRG collected from E13.5 embryos from pregnant mice carrying the conditional allele for the *Fxn* gene (*Fxn^L3/^*L3)(10) and enzymatically digested (35 min at 37°C in Trypsin 2.5‰) to dissociate the tissue. Cells (sensory neurons + glial cells) were seeded at a density of 7,500 cells/well in C-medium (and left incubated for 20 minutes (min) at room temperature (RT) before transferring the plates at 37°C, 5% CO_2_. The day after, cultures were treated with antimitotic (10 µM Uridine; 9.9 M FUdR) in NB-medium to remove proliferating glial cells from the culture (Fig. 1A, Fig. S1). NB-medium was then replaced every 2 or 3 days until the culture is treated.

**Figure 1:**
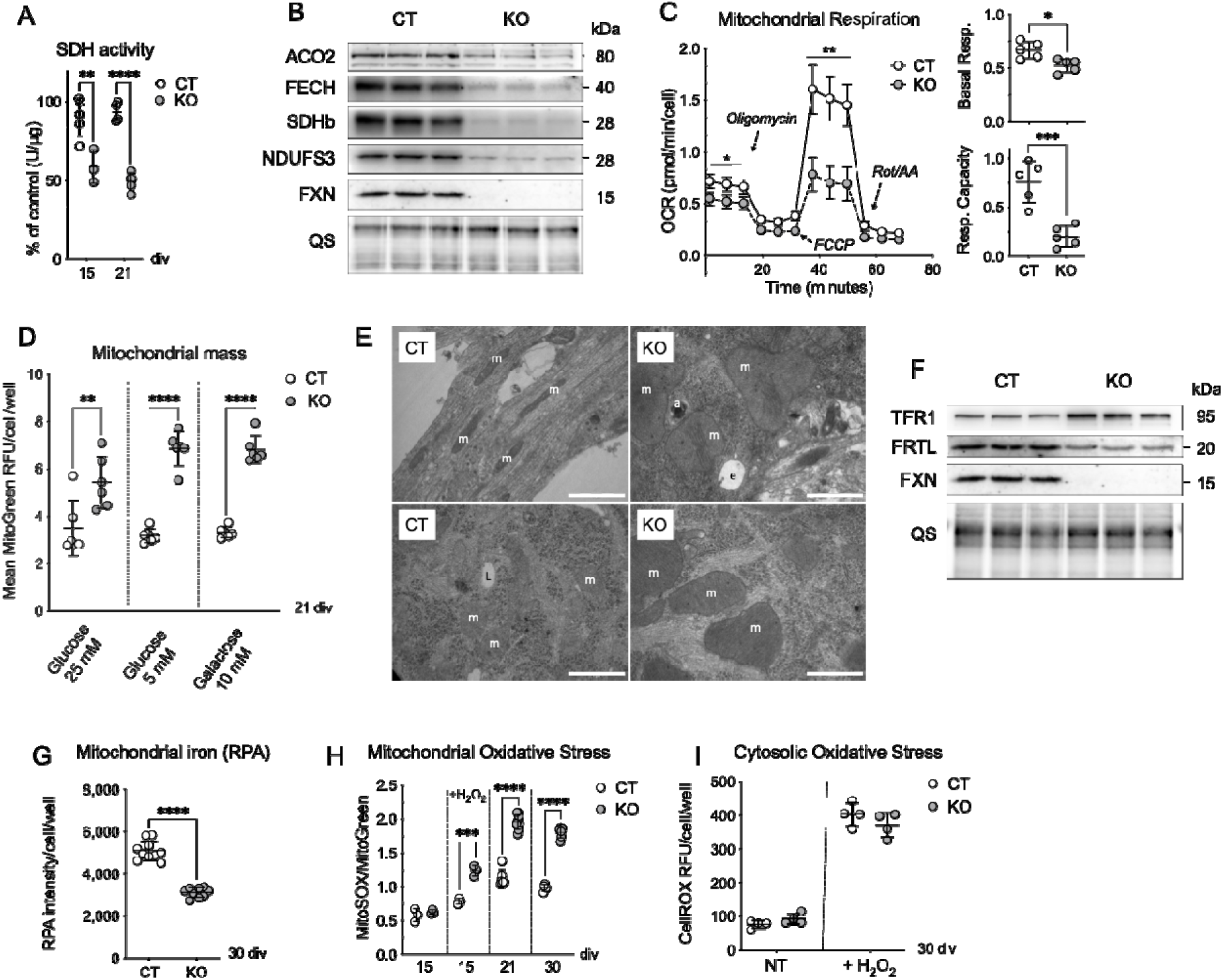
Sensory neurons depleted for FXN recapitulate several FA-associated biochemical and cellular pathological features *in vitro*. (A) Spectrophotometric measurements of succinate dehydrogenase (SDH) enzymatic activity on sensory neurons at 15 and 21 div. n = 5 control and 3-4 KO cultures. (B) Levels of mitochondrial proteins containing Fe-S clusters at 21 div. ACO2 = Aconitase; FECH = ferrochelatase; SDHb = Succinate Dehydrogenase B (Complex II); NDUFS3 = Complex I; FXN = FXN; QS=QuickStain (total protein levels). (C) Oxygen Consumption Rate (OCR) was measured by Seahorse Analyzer at 15 div through addition of different respiratory chain inhibitors, as indicated. Rot/AA = Rotenone/Antimycin A. Right panels show quantifications for basal respiration and maximal respiration capacity. n = 5 cultures each group. (D) Total mitochondrial mass measured in sensory neurons at 21 div by Mitotracker Green fluorescent probe under different culture conditions. Dots indicate averaged RFU (Relative Fluorescence Units) values from every cell of one well (96-well plates, about 300 cells/well). (E) Representative electron microscopy images of CT and KO sensory neurons after 30 div. m, mitochondria; e, endosome; a, autophagosome; L, lipids. Scale bars: 1 µm. (F) Protein levels of iron metabolism related proteins, transferrin receptor 1 (TFR1) and ferritin light chain (FRTL) in CT and KO sensory neurons at 21 div. (G) Mitochondrial iron content measured at 30 div by RPA sensor. RPA signal is inversely correlated to mitochondrial iron levels. n = 9-10 wells. (H) Mitochondrial oxidative stress assessed by MitoSOX fluorescent probe in sensory neurons at 15, 21 and 30 div, and normalized to mitochondrial content by Mitotracker Green FM. Cells at 15 div were treated with or without 10 µM H_2_O_2_. (I) Cytosolic oxidative stress assessed by CellROX. Fluorescent probe in sensory neurons at 30 div in DRG neurons, in the presence or absence of 10 µM H_2_O_2_ treatment. *p<0.05; **p<0.01; ***p<0.001; ****p<0.0001 (Student t-test). Error bars indicate SD.

Mice were maintained in a pathogen-free animal facility with a 12-hour light/dark cycle and provided with standard rodent chow (D04, SAFE, Villemoisson-sur-Orge, France), water and libitum. All animal procedures were performed in accordance with institutional animal care guidelines and approved by the local ethical committee (APAFIS# 54248-2024102512328818).

#### Plasmids

Bacterial LplA gene cDNA was cloned from *E. coli* (TOP10 strain) using the following primers: Fwd 5’-CACATTGCGCAAGAAATGCC3’ and Rev 5’ATGTCCACATTACGCCTGCT3’). A mitochondrial targeting sequence (MTS) was obtained from mouse *CoxIV*, added 5’ of the LplA cDNA fragment, and the entire mtLplA fragment was cloned into a pAAV-CMV vector (Stratagene). The AMPKα1-K56R-HA mutant was generated by overlap extension PCR. Two partially overlapping fragments were amplified independently and subsequently fused to introduce the point mutation.

PCR1 primers: Forward: 5′-attattATCGATATGCGCAGACTCAGTTCCTG-3′; Reverse: 5′-CCGGTTGAGTATCCTCACAG-3′. PCR2 primers: Forward: 5′-attattaagctttcaAGCGTAATCTGGAACATCGTATGGGTACTGTGCAAGAATTTTAATTAGATTTGC-3′; Reverse: 5′-GACATAAAGTGGCTGTGAGG-3′. The two PCR fragments shared a 35-nucleotide overlap, enabling fusion in a secondary PCR reaction. Both fragments were first amplified separately under standard conditions, purified, and then used as templates for the fusion PCR to yield the full-length AMPKα1-K56R-HA construct, which was cloned into a pAAV-CMV vector (Stratagene).

#### Viral infections

All viral vectors were produced and purified in the IGBMC in-house viral vector production facility (Strasbourg, France), following the helper-free method. For viral infections, AAV were diluted in NB medium (24) at a final concentration of 9×10^9^ vg/ml (unless indicated) and added to neuronal cultures overnight. The next day, AAV containing medium was fully replaced by fresh NB medium. Infections with either AAV9-empty (control) or AAV9-CreGFP (to excise exon 4 from the *Fxn* conditional allele) viruses were performed 3 hours after plating the cells (that is at 0 days in vitro, div). Infections using AAV9-CAG-hFXN-HA(17), AAV9-mtLplA and AAV9-AMPKα1-K56R-HA were performed at 7 div.

Doses of AAV9-CAG-hFXN-HA are as followed and expressed in VG/mL:

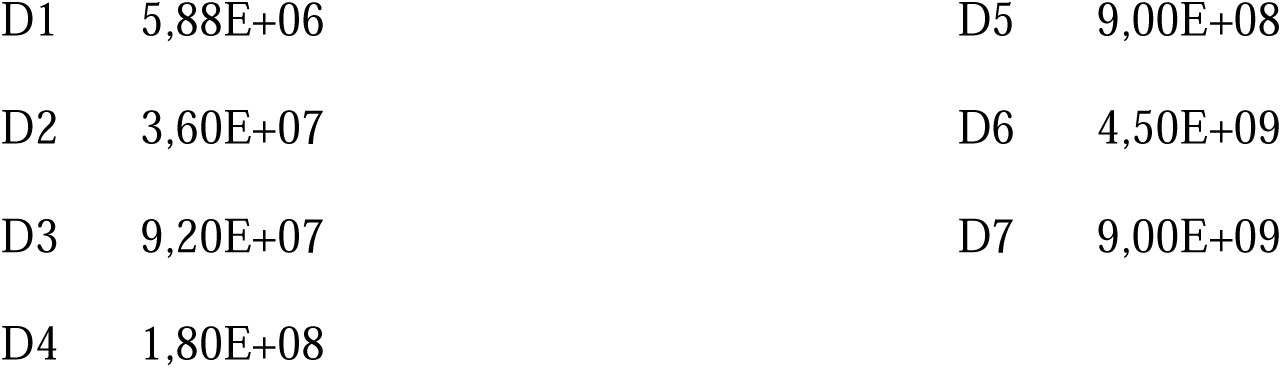

|  |  |  |  |
| --- | --- | --- | --- |
| D1 | 5,88E+06 | D5 | 9,00E+08 |
| D2 | 3,60E+07 | D6 | 4,50E+09 |
| D3 | 9,20E+07 | D7 | 9,00E+09 |
| D4 | 1,80E+08 |  |  |

#### Cellular treatments

Neuronal cultures were treated starting at 7 div (unless indicated) with either (±)-α-Lipoic acid (ALA; Merck T1395), AICAR (Merck, A9978), N-Acetyl-L-cysteine (Merck A7250), Compound 991 (Selleckchem ex229; S8654) or Rapamycin (Merck R8781). Fresh medium with chemicals was added every 3 to 4 days. In some experiments, cells were cultured either under normal (25 mM glucose), low glucose (5 mM) or glycolysis inhibitory (10 mM galactose in glucose-free NB-medium) conditions starting at 7 div.

#### Immunofluorescence experiments

DRG sensory neurons cultures were fixed for 15 min in 2% PFA diluted in PBSS and stained as described below. Lumbar mouse DRG were harvested from adult mice and fixed for 1h in 4% PFA. To obtain cryosection (8 µm) for immunofluorescence experiments, DRG were embedded in Shandon Cryomatrix resin (ThermoFisher 6769006) and snapfrozen in isopentane chilled on dry ice.

The following staining protocol was used for both sensory neuron cultures and DRG tissue floating sections. First, samples were permeabilized using 0.2% Triton X-100 diluted in PBSS, for 10 min, washed 3 times (5 min each) in PBSS, and blocked for 1 hour (h) with 1% BSA, 5% NGS, 0.1% Tween-20 in PBSS (PBSST). Samples were then incubated overnight at 4°C with the primary antibodies (Table S1) diluted in 1% BSA in PBSST. After 3 more washes in PBSST, secondary antibodies (Table S1) (1/2000) with DAPI (1/1000) were added and incubated for 30 min at RT. For actin staining, Rhodamine Phalloidin (1/2000 dilution, tebu-bio PHDR1) was added to the secondary antibody solution. Samples were finally washed twice in PBSST and twice in PBSS (5 min each) and mounted using Aqua-Poly/Mount (Polysciences18606) mounting medium.

#### Electronic microscopy

Sensory neurons cultures were fixed for 15 min in a solution containing 2.5% PFA, 2.5% glutaraldehyde, and 0.1M sodium cacodylate (pH = 7.2). Samples were then processed as previously described (10,25).

#### Protein extract and Western blots

Sensory neurons were collected and centrifuged at 6000 x g for 1 min. Pellets were resuspended in RIPA Lysis Buffer and incubated on ice for 20 min. After centrifugation for 10 min at 11.000 x g and at 4°C, supernatant was collected, and protein concentration measured by Bradford Assay. Desired quantity of total protein (generally 5µg/sample/gel) was pre-stained for 30 min using QuickStain Protein Labelling Kit (Amersham RPN4000) according to manufacturer’s instructions. Proteins were then mixed with loading buffer and boiled for 5 min, loaded on polyacrylamide gels, and separated by electrophoresis. Proteins were then tank-transferred onto a nitrocellulose membrane. After transfer, total protein fluorescence was acquired on an Amersham Imager 600 (GE Healthcare Life Sciences). Membranes were then blocked for 1h in Tris-buffered saline (TBS) containing 5% BSA and Tween-20 (TBST). Membranes were washed 3 times in TBST (10 min each) at RT and incubated with secondary antibodies (1/10.000 to 1/20.000) diluted in TBST for 45 min at RT. Membranes were washed 3 times in TBST (10 min each) and signal was revealed by SuperSignal™ West Femto Maximum Sensitivity Substrate (ThermoFisher 34096) using Amersham Imager 600.

#### Enzymatic activities

Succinate dehydrogenase (SDH) activity measurements were performed on 10 µg of proteins as previously described (10) on a Cary50 spectrophotometer (Varian SA). One unit of SDH activity is defined as the amount of SDH needed to generate 1.0 μmol of Dichlorophenolindophenol (DCPIP) per min (pH 7.2; 25°C). SDH activity was normalized to total protein content. Pyruvate dehydrogenase (PDH) and lactate dehydrogenase (LDH) activities were measured on 10 µg of proteins using colorimetric kits (Merck MAK183 and MAK066, respectively), following manufacturer’s instruction and using Synergy HTX reader (Agilent BioTek).

#### Seahorse analysis

Mitochondrial respiration was measured using a Seahorse Analyser XFe96 (Agilent) and Seahorse XF Cell Mito Stress Test Kit (Agilent 103015-100). Five thousand sensory neurons were seeded *per* well in compatible 96-well plates (Agilent 101085-004), and Seahorse analysis was performed at 15 div. Protocol was followed as per manufacturer’s instructions, with the following optimized concentrations of respiratory chain inhibitors: 2 µM Oligomycin, 0.5 µM FCCP, and 0.5 µM of Rotenone/Antimycin A. Mixing steps (Wave Controller 2.4) were reduced to 20 seconds to prevent detaching of the cells from the bottom of the well.

#### Survival, soma area, mitochondrial mass, MitoSOX, CellROX and TMRM measurements

Sensory neurons were seeded in black 96-well plates (Greiner 655090) and were stained for 3h (at 37°C, 5% CO_2_) with Hoechst 33342 (1 mg/ml; Merck 94403), Mitotracker Green FM (50 nM; ThermoFisher M7514), and/or MitoSOX (2.5 µM; ThermoFisher M36008), or CellROX (5 µM; ThermoFisher C10422) or TMRM (200 nM; Invitrogen I34361) diluted in NB culture medium the day of the analysis at different time points depending on the experiment. To determine the sensitivity to exogeneous oxidative stress, oxidative treatment was performed with 10 µM H_2_O_2_ incubation for 30 min at 37°C, 5% CO_2_ before MitoSOX or CellROX staining. Cells were then washed twice in PBSS (10 min each), fixed in 2% PFA for 15 min (except for TMRM, acquired in live cells), and washed twice with PBSS (10 min each). Fluorescence was acquired on a high content reader (Thermofisher Cellomics Arrayscan XTi) and images were analyzed by the Cellomics HCS Studio software. Each dot indicates average fluorescence from one well in a 96-well plate (about 300 cells/well). Hoechst segmentation was also used to count cells for survival analysis. In addition to labeling nuclei, Hoechst provided enough signal in the soma to allow segmentation and measurement of soma area. Pvalb-positive soma area of Pvalb-Cre mouse (FxnL3/L-; Pvalbtm1(Cre)Arbr/J, (17)) were measured using ImageJ. Signal intensity, and area of each sensory neuron was acquired independently, through cell segmentation.

#### Mitochondrial iron content

Mitochondrial iron was measured in sensory neurons seeded in black 96-well plates, using the fluorescent non-toxic iron sensor RPA (Axxora SQX-RPA). Sensory neurons were stained with Hoechst 33342 (1 ug/mL; Merck 94403) and 2.5 µM RPA in NB medium for 10 min at 37°C and washed three times in PBSS. In conditions of iron supplementation, cells were pretreated for 3 days with 100 µM Ferric Ammonium Citrate (FAC, F5879, Sigma) in NB medium. In conditions of iron chelation, cells were pretreated for 20 minutes with 2 mM of 1,10-phenantroline (P9375, Sigma) in NB medium. Fluorescence was acquired on a high content reader (Thermofisher Cellomics Arrayscan XTi), and images were analyzed by the Cellomics HCS Studio software. Signal intensity of every cell was acquired independently through cell segmentation.

#### ATP measurements

ATP levels were measured using a fluorometric kit (Merck MAK190), following manufacturer’s instructions. Amount of total protein used was optimized to 10 µg/sample/well.

#### pH measurements

pH in culture media was indirectly assessed by measuring absorbance of phenol red at 415 nm and 560 nm. Ratiometric analysis 415nm/560nm was calculated to obtain pH (26).

#### Reverse transcription and real time quantitative PCR (RT-qPCR)

Total RNA was extracted from frozen sensory neuron cultures pellets using TRIzol reagent (ThermoFisher 15596026), according to the manufacturer’s protocol, and treated with DNAse I (Roche 4716728001). cDNA was generated by reverse transcriptase using the SuperScript IV Kit (Invitrogen 18091050), following the manufacturer’s protocol. Quantitative RT-PCR was performed using the SYBR Green I Master mix (Roche 04887352001) on Light Cycler 480 (Roche) with primers listed in Table S3. Gene expression was normalized to GAPDH expression, and then in percent of CT.

#### Amino acid measurements

Amino acid levels in sensory neurons cultures were measured using LC-MS/MS chromatography (LC-MS/MS Xevo TQS micro; Waters) and the Amino Acid Analysis (AAA) kit (Chromsystems) in collaboration with the lab of Dr. Daniel Brumaru (Hôpital de Hautepierre, Strasbourg). Samples were lysed in a hypotonic buffer (10 mM KH_2_PO_4_, 2 mM EDTA, pH 7.8) and the measures were performed on according to the manufacturer’s protocol. Amino acid levels were normalized to total protein levels.

#### Statistical analyses

All data are presented as mean ± SD. Statistical analyses were carried out using GraphPad Prism software (La Jolla, USA). Student’s t tests were used to compare two groups, one-way ANOVA (Kruskal-Wallis test) was used to compare 3 or more groups. Two-way ANOVA with Geisser Greenhouse Correction was used to compare size combined to genotypes and classes of cell area/time points. A value of p < 0.05 was considered as significant. Replicates (n) were considered as biological replicates as they correspond to independent cultures issued from a pool of mouse embryos, independently infected and independently grown until analysis. Fluorescent and blot images show representative results of several experiments. For cell area and fluorescence experiments performed in 96 well plates, graph dots indicate averaged result from every cell of one well (about 300 cells/well).

## Results

### FXN-depleted DRG sensory neurons recapitulate FA molecular and cellular features

To investigate the impact of FXN deficiency in a disease-relevant cell type, we previously established an *in vitro* culture model for FA based on mouse primary DRG sensory neurons carrying the conditional *Fxn* allele (24). DRG were harvested at embryonic stage E13.5 and after dissociation and seeding, the cells were infected with an AAV9-CreGFP vector (KO) or with an empty AAV vector as a control (CT) (Fig. S1A). Glial proliferation was subsequently suppressed using antimitotic treatment, resulting in highly enriched neuronal cultures (Fig. S1B). Infection with the AAV9-CreGFP virus to excise the conditional *Fxn* allele, resulted in near-complete depletion of FXN expression levels in the culture after 7 div (Fig. S1C), in accordance with the >95% viral infection efficiency (24). We next characterized the neuronal composition of the cultures using sensory neuron subtypes-specific markers. A large proportion of neurons expressed NF200 (marker for large myelinated sensory neurons, mechanoreceptive and proprioceptive), parvalbumin (Aß-LTMRs and proprioceptive neurons), and TrkC (mechanoreceptive and proprioceptive neurons), consistent with enrichment in large sensory neuron subtypes. In addition, a substantial proportion of cells was positive for the nociceptive specific marker calcitonin gene-related peptide (CGRP) (Fig. S1 D). Moreover, expression of these markers was stable over time (7 to 25 div (Fig. S1 D), with no major differences between control (CT) and FXN-deficient (KO) cultures (Fig. S1 D, E).

We then assessed key biochemical and cellular consequence of frataxin depletion (Fig. 1). Activity of the Fe-S enzyme succinate dehydrogenase (SDH) was significantly reduced in KO sensory neurons as early as 15 div and further decrease at 21 div (Fig. 1A). In addition, a substantial decrease in mitochondrial and cytosolic Fe-S cluster containing proteins was observed by western blot at 21 div (Figs. 1B and S2A), in line with previously reported instability of the apo-proteins or complexes in the absence of Fe-S cluster (25,27).

Mitochondrial function was strongly impaired, as shown by Seahorse analysis revealing reduced basal respiration rate and a severe reduction of respiratory capacity in KO neurons compared to the CT at 15 div (Fig. 1C). This was accompanied by evidence of metabolic reprogramming (28), including increased glycolytic activity, as indicated by medium acidification (Fig. S2B) and increased activity of lactate dehydrogenase (Fig. S2C). Mitochondrial mass was found increased in KO cells which was exacerbated under low glucose or glycolysis inhibiting (galactose) conditions (Fig. 1D). Accordingly, several mitochondrial proteins as well as transcriptional factors involved in mitochondrial biogenesis were found to be increased at the translational levels (Fig. S2D). At the ultrastructural level, KO cells contained giant mitochondria with abnormal structures including collapsed cristae (Fig. 1E).

A direct consequence of Fe-S deficit is the dysregulation of iron metabolism through the Fe-S-dependent activation of the Iron Responsive Element (IRE)-binding Protein IRP1 (29). An increase of transferrin receptor 1 (TFR1) and a decrease of ferritin light chain (FRTL) was observed in KO cells at 21 div (Fig. 1F), consistent with an activation of IRP1. In contrast, no significant changes in TFR1 and FRTL were detected at 15 dpi (Fig. S2E), indicating that alterations of the iron homeostasis occur after the onset of Fe-S protein deficiency. Increased mitochondrial iron content is another hallmark of the FA disease. Mitochondrial iron content was measured using the fluorescent mitochondrial iron indicator RPA, which signal depends on labile iron content (30). RPA signal was found to be decreased in the KO sensory neurons at 30 div compared to CT sensory neurons (Fig. 1G), consistent with increased labile mitochondrial iron levels and/or altered mitochondrial membrane potential. Importantly, iron chelation increased RPA fluorescence whereas iron loading decreased it in both CT and KO neurons, confirming that the probe remains responsive to mitochondrial iron in both conditions (Fig. S2F).

Mitochondrial electron transport chain dysfunction is commonly associated with elevated production of reactive oxygen species (ROS)(31). Numerous FA models present an increased sensitivity to oxidative stress(11). To assess mitochondrial oxidative stress, we used the mitochondrial-specific fluorescent probe MitoSOX. At early time point (15 div), KO sensory neurons displayed an increase in sensitivity to exogenous oxidative stress compared to CT, without detectable increase in basal mitochondrial ROS levels (Fig. 1H). At later stages (21 and 30 div), KO sensory neurons exhibited an increase in endogenous mitochondrial oxidative stress relative to CT neurons (Fig. 1H). In contrast, measurements using the cytosolic ROS probe CellROX at 30 div revealed no increase in cytosolic oxidative stress under basal condition and in the presence of exogenous stress in KO sensory neurons compared to CT (Fig. 1I).

Altogether, these findings demonstrate that this model reproduces several key biochemical and cellular features of frataxin deficiency in sensory neurons.

### Frataxin depletion impairs soma size without affecting the survival of sensory neurons

To assess whether frataxin deficiency affects neuronal viability, cell survival was assessed over time. No significant difference in cell number was observed between KO and CT sensory neurons, even at late stages (30 div) (Fig. 2A). Furthermore, KO neuron viability was maintain under glycolysis-inhibiting conditions (Fig. S3). In contrast, a significant difference in soma growth dynamics between KO and CT sensory neurons was observed. From 7 to 15 div, KO and CT sensory neurons exhibited comparable soma growth (Fig. 2B). While CT soma continued to grow until 21 div, KO soma size plateaued, and subsequently declined slightly by 30 div (Fig. 2B). Quantitatively, KO somas were 1.3-fold smaller than CT at 21 div, and this difference increased to 1.5-fold by 30 div (Fig. 2B). This reduction was confirmed by 3D imaging, which showed a global decrease in KO neuron soma volume compared to CT (Fig 2C). Analysis at the single-cell level revealed a reduced proportion of large soma size neurons in the KO population (Fig. 2D). Since total cell number remains unchanged, these results suggest a global shift in cell size distribution rather than selective loss of large neurons.

**Figure 2:**
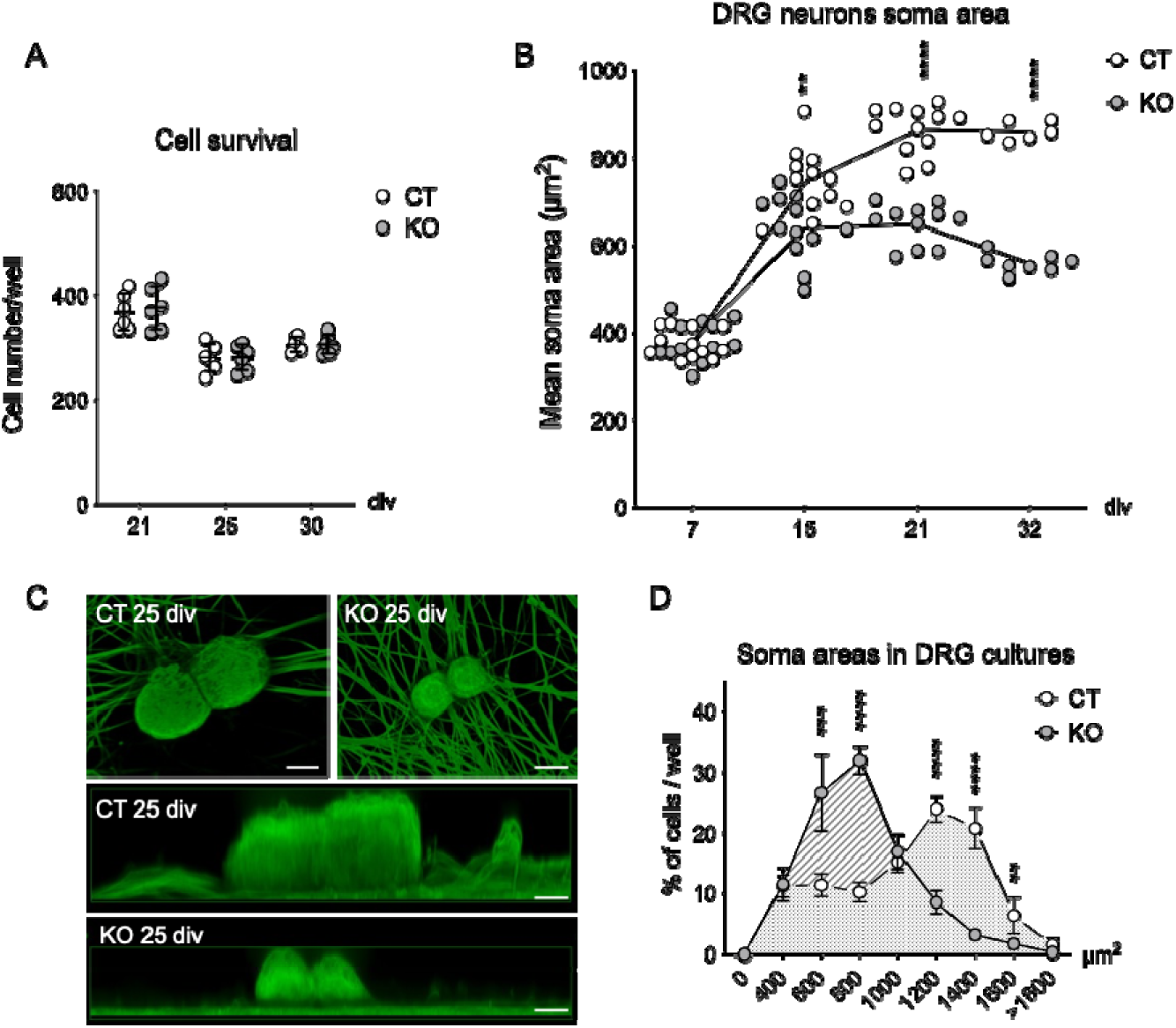
Sensory neurons present similar survival rate but decreased soma area in the absence of FXN. (A) Cell count over time measured by Hoechst staining. n = 6 wells (B) Sensory neuron soma area measured by Hoechst staining (reflection of the signal throughout the cell body) at different time points. Dots indicate averaged areas from about 300 cells in one well (8-10 wells analyzed)). n = 6-12 independent cultures. (C) 3D representation of NF200-positive cells in CT and KO cultures at 25 div, acquired by z-scan with confocal microscopy. Scale bar = 10 µm. (D) Distribution frequency plots of soma areas in sensory neuron cultures. n = 6 wells (about 300 cells/well) at 25 div. (B, D) Two-way ANOVA with Geisser Greenhouse Correction; **p<0.01; ***p<0.001; ****p<0.0001. Error bars indicate SD.

### Soma size is rescued by frataxin in a dose-dependent manner

To assess whether the KO phenotype could be rescued in a frataxin dose-dependent level, DRG neurons were transduced with increasing doses of an AAV vector encoding HA-tagged human frataxin (AAV9-CAG-hFXN-HA (17)). Anti-HA immunostaining confirmed the dose-dependent transgene expression (Fig. S4). Interestingly, several hallmarks of the FA-phenotype, including Fe-S deficit (assessed indirectly via protein stability; Fig. 3A), mitochondrial oxidative stress (Fig. 3B), and mitochondrial membrane potential impairment (Fig. 3C), were progressively corrected as frataxin expression increased. Importantly, increasing frataxin levels also restored soma size in KO neurons in a dose-dependent manner (Fig. 3D).

**Figure 3:**
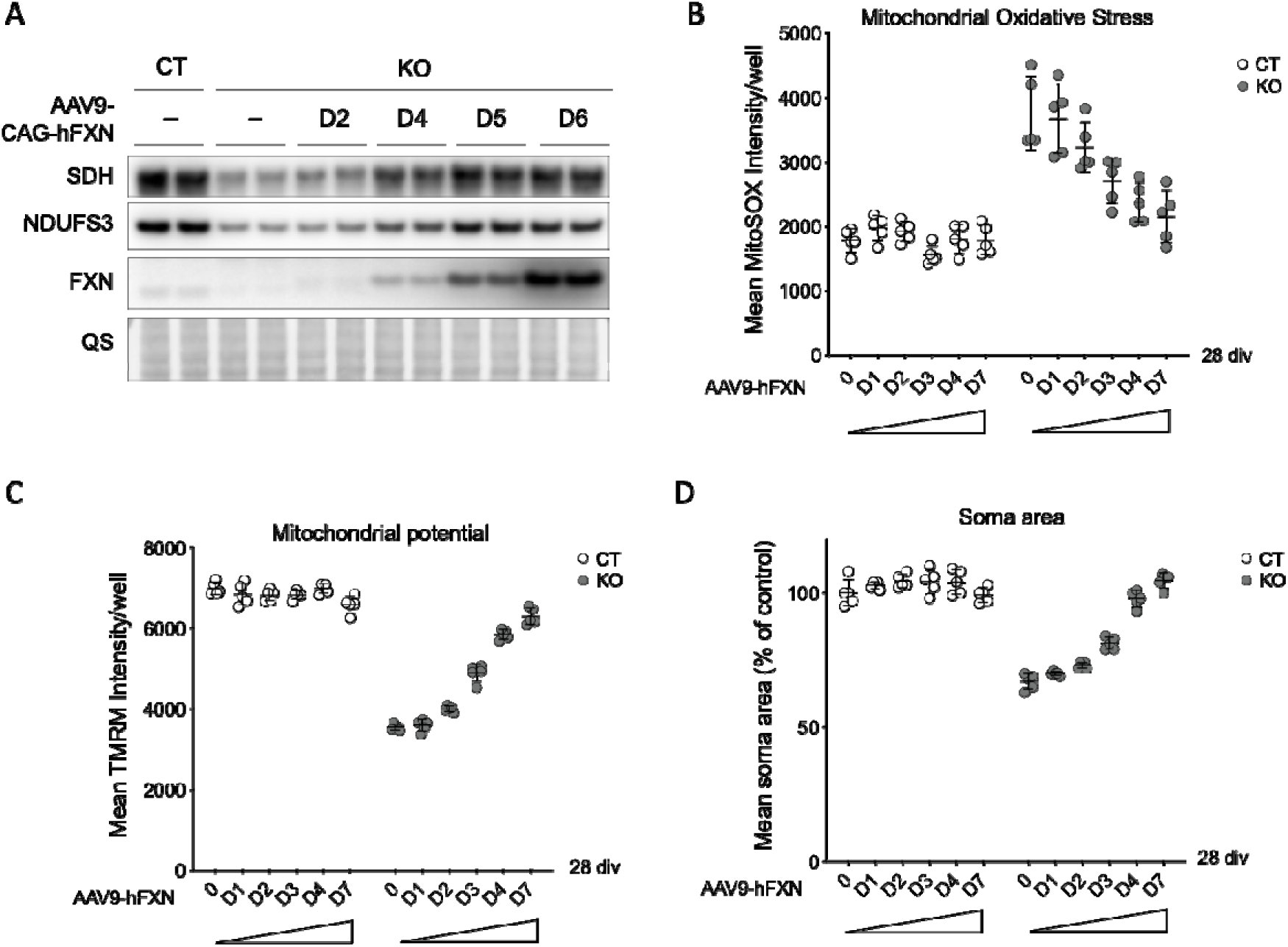
Dose-dependent reversal of phenotype in KO sensory neurons by exogenous hFXN expression. Control (CT) and FXN-depleted (KO) sensory neurons were cultured for 7 div, infected with different concentrations of AAV-CAG-hFXN-HA viral particles. (A) Levels of the Fe-S containing proteins SDH and NDUFS3 in CT and KO sensory neuron cultures without or with increasing doses of AAV-CAG-hFXN-HA infection: 3.6 x10^7^ vg/mL (D2); 1.8 ×10^8^ vg/mL (D4); 9 ×10^8^ vg/mL (D5); 4.5 ×10^9^ vg/mL (D6). Note, anti-FXN antibody (clone 4F9) has an extensively higher affinity for hFXN than mouse FXN (mFXN). (B) Mitochondrial oxidative stress assessed by MitoSOX signal following increase in hFXN expression. (C) Mitochondrial membrane potential measured by TMRM following increase in exogenous hFXN expression. (B, C). Dots indicate averaged RFU from every cell of one well (96-well plates, about 300 cells/well). n = 4 wells. (D) Measurements of sensory neuron soma area after increase in hFXN expression (28 div). Dots indicate averaged areas from every cell of one well (96-well plates, about 300 cells/well). n = 4 wells. Two-way ANOVA with Geisser Greenhouse Correction was applied in B, C and D. ***p<0.001; ****p<0.0001. Error bars indicate SD.

### Lipoic acid-dependent pathways are impaired in frataxin-deficient neurons

As lipoic acid (LA) synthesis depends on the Fe-S cluster-dependent enzyme lipoic acid synthase (LIAS), we next evaluated protein lipoylation status in KO neurons. Quantitative analysis revealed a pronounced decrease in LA conjugation to both PDH and α-ketoglutarate dehydrogenase (KGDH), two key enzymes of the Krebs cycle (Fig. 4A). Importantly, PDH E2 subunit (PDC-E2) protein levels remained unchanged, suggesting that the reduction in lipoylation reflects functional impairment of LIAS activity. Consistent with this, LIAS protein levels were significantly reduced in KO sensory neurons (Fig. 4A), likely secondary to destabilization resulting from defective Fe-S cluster biogenesis. A significant reduction (∼ 2.6-fold) in pyruvate dehydrogenase (PDH) activity, which requires covalent attachment of LA to PDC-E2, was detected in KO sensory neurons, indicating compromised mitochondrial metabolism (Fig. 4B).

**Figure 4:**
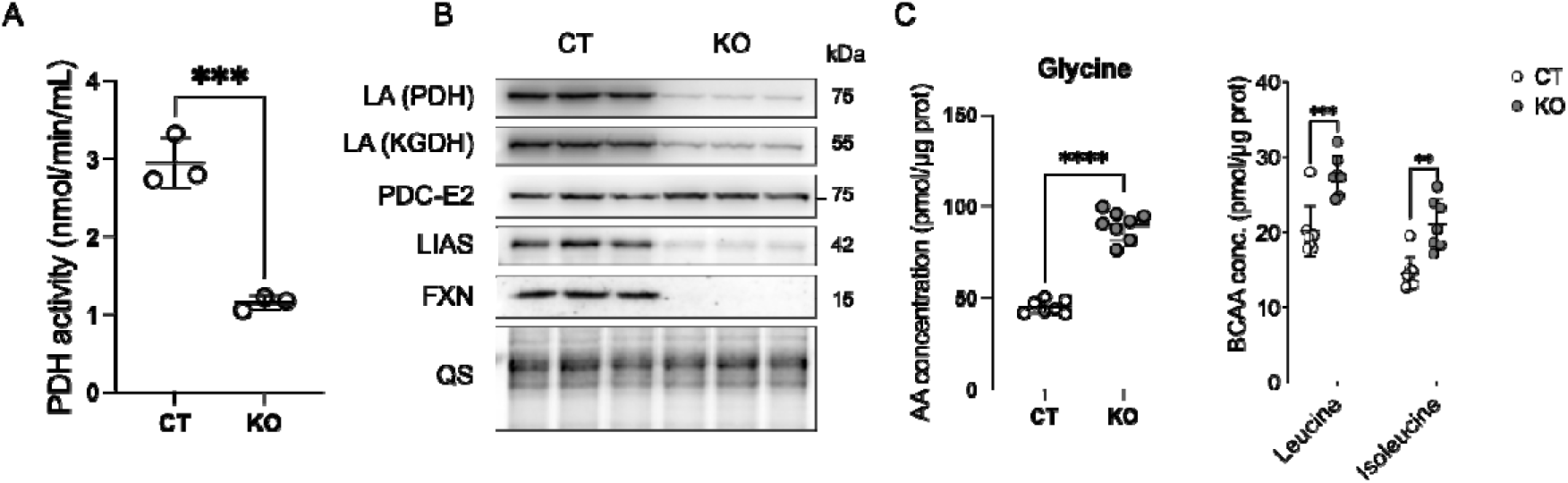
Lipoic acid-associated pathways are affected in frataxin deficient sensory neurons. (A) PDH activity of CT and KO cells at 21 div. (B) Western Blot of Lipoic Acid (LA) on recipient enzymes PDH and KGDH, total PDC-E2, LIAS and FXN in CT and KO cells at 21 div. QS staining used as a loading control for quantification. (C) Measure of concentration of glycine, leucine and isoleucine in CT and KO cells at 30 div. (A, C) One-way ANOVA followed by Tukey for multiple comparison. *p<0.05; **p<0.01; ***p<0.001; ****p<0.0001. Error bars indicate SD.

Since LA is also a required cofactor for mitochondrial degradation pathways of glycine, lysine, and branched-chain amino acids (BCAAs), we investigated the metabolic consequences of impaired LA-dependent enzyme function. Elevated levels of glycine and BCAAs were detected in KO neurons at 30 div (Fig. 4C), consistent with defective amino acid catabolism due to insufficient lipoylation of dehydrogenase complexes.

To restore LA-dependent enzyme activity in KO neurons, we expressed of Lipoate ligase A (LplA), a bacterial enzyme from the salvage pathway capable of attaching exogenous LA to target proteins, a function absent in mammalian systems, which rely exclusively on *de novo* LA synthesis (32). For mitochondria targeting, LplA was fused to the human COXV mitochondrial targeting sequence and delivered via AAV9-mediated transduction (mtLplA). Immunofluorescence indicated partial mitochondrial localization of mtLplA, likely due to saturation of the mitochondrial import machinery under conditions of high expression, consistent with the presence of both processed and unprocessed forms detected by western blot (Fig. S5A,B). Expression of mtLplA partially restored LA loading to PDH and KGDH in the absence of exogenous LA, and fully restored lipoylation upon supplementation with ≥ 10 µM α-lipoic acid (ALA) (Fig. S5B). Correspondingly, PDH enzymatic activity was fully rescued under these conditions (Fig. S5C).

Interestingly, while mtLplA expression restored biochemical defects of LA loading and PDH activity, it failed to normalize soma size phenotype (Fig. S5D). Phenotypic rescue required high-dose ALA supplementation, which induced a dose-dependent increase in soma size and shifted the KO soma size distribution toward larger neurons (Fig. S5E). Although mtLplA expression increased the proportion of large neurons, ALA, either alone or combined with mtLplA, resulted in a broader rescue across the neuronal population. These observations suggest that ALA may have additional lipoylated proteins-independent actions, possibly related to its antioxidant properties, that contribute to the restoration of neuronal soma size.

### The AMPK pathway is activated in frataxin-deficient neurons

The AMP-activated protein kinase (AMPK) signaling pathway plays a key role in regulating cell growth, metabolism and autophagy (33). AMPK senses cellular energy status by responding to fluctuations in cellular ATP/AMP ratio. Given the strong decrease in mitochondrial function resulting in low ATP levels (Fig. 5A), we hypothesized that AMPK pathway could contribute to the growth defects observed in KO neurons. Western blot analysis revealed a clear activation of the AMPK pathway at 21 div in KO sensory neurons (Fig. 5B). Specifically, the phospho-AMPKα-Thr172 activation mark was increased by 1.7-fold relative to controls (Fig 5B). In parallel, phosphorylation of several downstream AMPK targets, including ACC, ULK1 and Raptor, were significantly increased in KO neurons (Fig. 5B). Notably, the phosphorylation of ULK1, an autophagy regulator, at Ser555 was increased by 2.4-fold (Fig. 5B), accompanied by elevated phosphorylation of its downstream target Beclin-1 at Ser15 (Fig. S6A).

**Figure 5:**
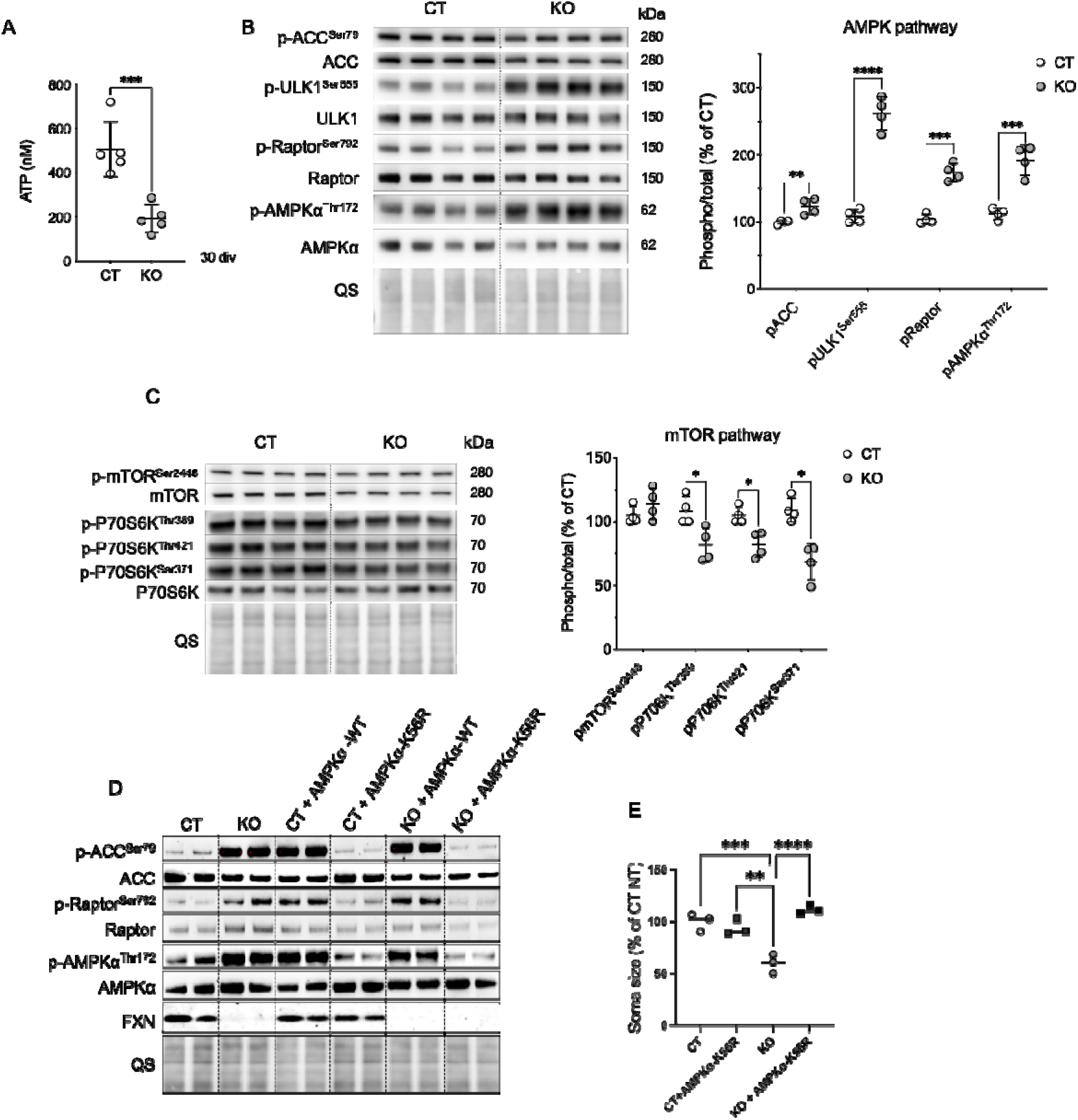
Activation and Rescue of AMPK/mTOR pathways in sensory neurons depleted for FXN. (A) ATP content in CT and KO sensory neurons at 30 div. (B) Western blot analysis of AMPK pathway in CT and KO cells. Right panel indicates quantification of the Western Blot. QS staining used as a loading control for quantification. (C) Western Blot analysis of the mTOR pathway. Right panel indicates quantification of the Western Blot. QS staining used as a loading control for quantification. (D) Western blot analysis of AMPK pathway in CT and KO cells with or without overexpression of a dominant-negative AMPK_α_ mutant (AMPK_α_^K56R^) or AMPK_α_ WT. (E) DRG neuron soma area measured in CT and KO cells with or without AMPK_α_^K56R^ expression. Areas are expressed in percentage of averaged CT. *p<0.05; **p<0.01; ***p<0.001; ****p<0.0001. Error bars indicate SD. Student t-test was applied in A, B, C and one-way ANOVA followed by Tukey for multiple comparison was used in E.

AMPK is known to inhibit cell growth partly through suppression of the mammalian target of rapamycin complex 1 (mTORC1) pathway (34). mTOR controls cell growth by promoting anabolic processes, such as protein, lipid and nucleotide synthesis, and suppressing catabolic processes, such as autophagy (34). AMPK directly phosphorylates Raptor, a regulatory associated protein of mTOR, thereby blocking mTORC1 activity(33). Consistent with this, phosphorylation of Raptor at Ser792 was increased by 1.7-fold in the KO neurons (Fig. 5B). Furthermore, phosphorylation of mTOR downstream target P70S6K was significantly decreased on several residues (Fig. 5C), indicating inhibition of the mTOR/S6K axis. Interestingly, pharmacological activation of the AMPK pathway with AICAR or Compound 991 treatments reduced soma size in CT sensory neurons (Fig. S6B-D). Similarly, pharmacological inhibition of mTOR pathway using Rapamycin also led to decrease soma size in CT sensory neurons (83 ± 2%) (Fig S6B, S6E). Restoration of frataxin expression in KO neurons progressively normalized both AMPK activation and mTOR pathways inhibition (Fig. S7A).

To directly test whether AMPK activation drives soma size reduction, we inhibited endogenous AMPK activity by overexpressing a dominant-negative AMPKα mutant (AMPKα^K56R^). This approach effectively suppressed AMPK activation, as shown by reduced phosphorylation of AMPKα at Thr172 and decrease phosphorylation of its downstream targets ACC and Raptor (Fig. 5D). Importantly, inhibition of AMPK by AMPKα^K56R^ led to a significant increase in soma size in KO neurons, restoring it to control levels (Fig. 5E). These finding indicate that AMPK activation directly contributes to the soma size phenotype observed in frataxin-deficient neurons.

### Antioxidant properties of ALA contribute to phenotypic rescue and prevents AMPK activation in frataxin-deficient neurons

As previously seen, treatment with exogenous LA in the form of ALA partially rescued soma size defects in KO neurons in a dose-dependent manner (Fig. 6A). ALA treatment also improved protein lipoylation, as evidence by a partial restoration of LA conjugation to PDC-E2 (13 ± 2% increase; Fig. S8A). However, ALA had minimal impact on glycine and BCAA levels, although a trend toward normalization was observed (Fig. S8B). ALA treatment significantly reduced mitochondrial oxidative stress in the KO neurons (Fig 6B). The dose-dependent rescue of soma size by ALA prompted us to examine whether this effect might be mediated via AMPK signalling. Phosphorylation of AMPKα at Thr172, a key marker of AMPK activation, was strongly decreased following ALA treatment in KO neurons (Fig 6C). In parallel, phosphorylation of downstream AMPK targets, including ULK1, was also significantly reduced (Fig 6C), indicating suppression of AMPK activity. ALA also restored intracellular ATP levels in KO neurons (Fig 6D), suggesting an improvement in cellular bioenergetics. Together, these findings indicate that ALA treatment is associated with improved metabolic and redox parameters and reduced AMPK activation, which correlates with the rescue of neuronal growth.

**Figure 6:**
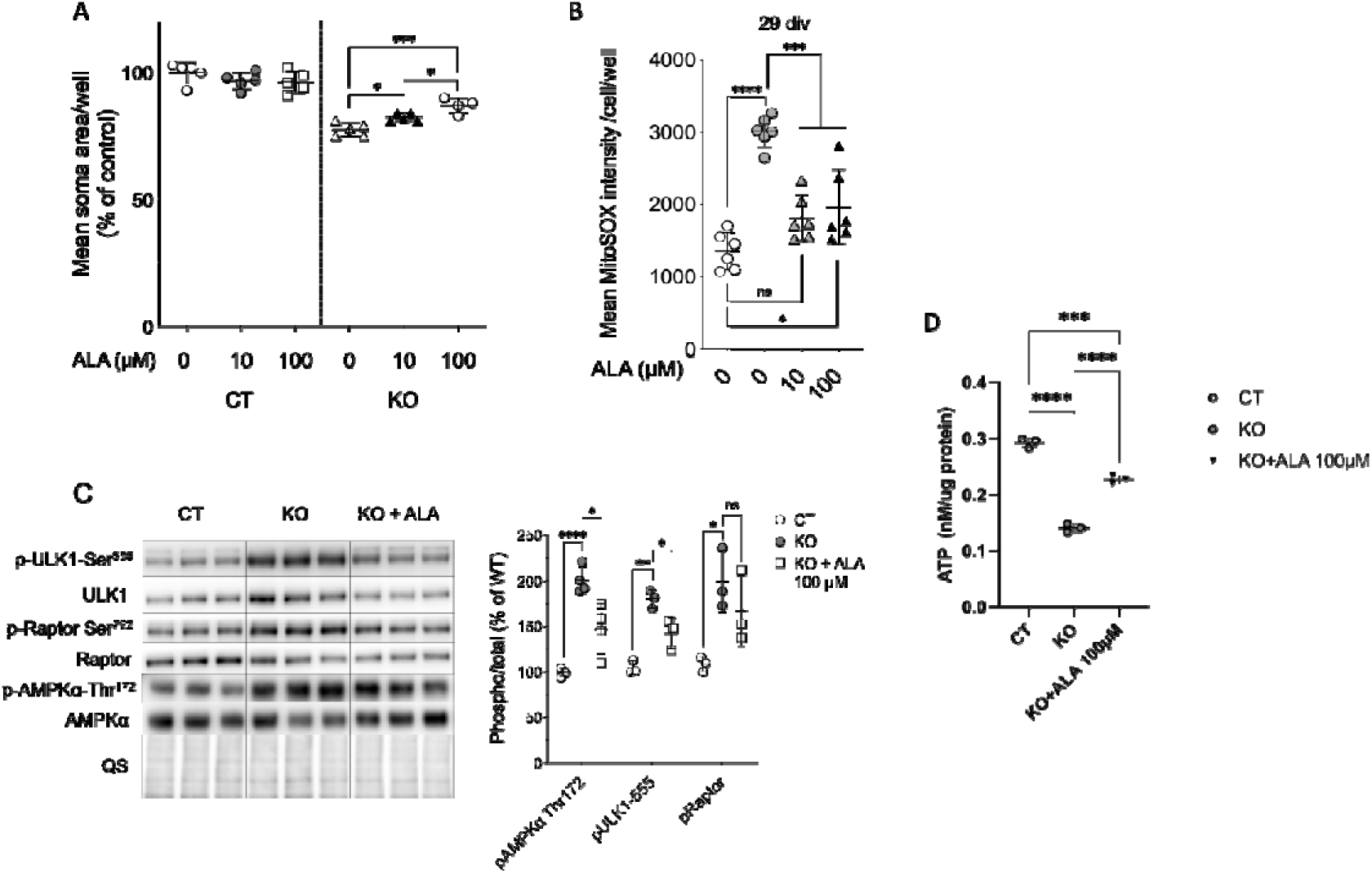
Antioxidant properties of ALA contribute to phenotypic rescue and prevents AMPK activation in frataxin-deficient neurons. (A) DRG neuron soma area measured in 96-well plates at 21 div, following ALA treatment. Dots indicate averaged areas from every cell of one well (96-well plates, about 300 cells/well). n = 4-5 wells. (B) Mitochondrial oxidative stress assessed by MitoSOX. MitoSOX fluorescence was acquired at 29 div in DRG CT or KO neurons treated with ALA. Dots indicate averaged values from every cell of one well (96-well plates, about 300 cells/well). (C) Western Blot of phosphorylated and total proteins of AMPK pathway, In CT or KO neurons treated with ALA (100 µM). Right panel indicates quantification of the Western Blot. QS staining used as a loading control for quantification. (D) ATP content in CT and KO sensory neurons treated or not with ALA at 30 div. (A-D) One-way ANOVA followed by Tukey for multiple comparison. *p<0.05; **p<0.01; ***p<0.001; ****p<0.0001. Error bars indicate SD.

## Discussion

In this study, we established a primary neuronal model of FA based on mouse DRG sensory neurons with complete FXN depletion. This model reproduces key biochemical and cellular features associated with frataxin deficiency, including early and progressive loss of Fe-S proteins, dysregulated iron metabolism, and severe mitochondrial dysfunction characterized by decreased mitochondrial oxidative respiration leading to energy deficit, increased mitochondrial mass, and oxidative stress. These phenotypes are consistent with those observed in other FA-relevant cell types such as cardiomyocytes (10,35,36).

Importantly, our data support a model in which full FXN depletion first induces a deficit of Fe-S synthesis, subsequently triggering downstream consequences such as iron dysregulation, mitochondrial proliferation, and redox imbalance. Notably, markers of iron homeostasis remained unchanged at early time points, indication that Fe-S deficiency precedes detectable dysregulation of iron metabolism in this system. Mitochondrial oxidative stress emerged progressively and was largely restricted to the mitochondrial compartment, consistent with a primary mitochondrial origin of the redox imbalance. These observations are in line with findings from DRG sensory neurons from a GAA-based FA mouse model (37).

Mitochondrial mass was increased in FXN-deficient neurons. Since this was not accompanied by improved function, we interpret it as a compensatory response to bioenergetic stress or as a result of impaired mitophagic clearance of damaged organelles. Although mitophagy flux was not directly evaluated, the presence of abnormally enlarged mitochondria with collapsed cristae strongly suggests impaired mitochondrial quality control, a hypothesis that warrants further investigation. Notably, similar abnormalities were also observed in other models (17,38), reinforcing the relevance of this phenotype.

Interestingly, despite pronounced mitochondrial dysfunction, FXN-deficient DRG neurons remained viable for several weeks *in vitro*, even under glucose-restrictive conditions. This metabolic resilience contrasts with the rapid death of dividing cells following FXN loss (39,40), likely due to the critical dependence of cell division and DNA replication on Fe-S proteins essential for these processes (41,42). These findings are consistent with observations from the *Pvalb* conditional knockout mouse model, in which proprioceptive neurons survived for several weeks complete FXN depletion *in vivo* (17). This suggests that, although functionally compromised, sensory neurons retain sufficient plasticity to withstand FXN loss over extended periods, providing a potential therapeutic window before irreversible degeneration occurs, as evidenced by full post-symptomatic rescue observed in gene therapy studies (17,25).

A key finding of this study is the identification of a defect in neuronal soma growth in FXN-deficient neurons. Rather than inducing cell death, FXN depletion impaired the ability of sensory neurons to achieve and maintain normal soma growth, resulting in a global shift in cell size distribution. Importantly, re-expression of human frataxin rescued soma size in a dose-dependent manner, strongly indicating that the growth deficit is a direct and frataxin-specific phenotype. While reduced neuronal size has not been extensively characterized at the cellular level in patient tissues, neuropathological studies consistently report hypoplasia and atrophy of the DRG and spinal cord in FA, suggesting that impaired neuronal growth and maturation may contribute to disease pathology. In this context, our findings support the idea that frataxin deficiency may affect developmental processes in addition to driving neurodegeneration.

FXN-deficient neurons exhibited substantial disruption of lipoic acid (LA)-dependent pathways. Lipoylation of both PDH and KDGH was impaired. Since LA biosynthesis depends on the Fe-S enzyme lipoid acid synthase (LIAS), these findings are consistent with compromised LIAS activity secondary to Fe-S depletion. Protein levels of LIAS were also reduced, likely due to destabilization of the apoenzyme. Consequently, key metabolic pathways relying on lipoylated enzymes, including the Krebs cycle and the mitochondrial degradation of glycine and branched amino acids (BCAAs), were affected with PDH activity significantly reduced and elevated levels of glycine and BCAAs in FXN-deficient neurons. To bypass the endogenous LA synthesis defect, we expressed a mitochondria-targeted version of the bacterial enzyme lipoate ligase A (mtLplA), which salvaging exogenous LA by attaching it to recipient proteins. While mtLplA expression restored PDH and KGDH lipoylation and PDH activity, it failed to rescue the soma size defect. In contrast, high-dose α-lipoic acid ALA supplementation induced a robust, dose-dependent rescue of soma size, both alone or in combination with mtLplA. Considering that ALA alone does not rescue PDH activity, these findings suggests that ALA exerts effects beyond its role as a cofactor, likely due to its antioxidant properties and impact on cellular signaling.

Mechanistically, we identified AMP-activated protein kinase (AMPK) as a central mediator of the growth defect. FXN-deficient neurons exhibited severe mitochondrial dysfunction, reflected by reduced respiratory activity and ATP levels, alongside elevated mitochondrial oxidative stress. This bioenergetic and redox imbalance provides two synergistic signals known to activate AMPK: a high AMP/ATP ratio and accumulation of reactive oxygen and nitrogen species (ROS/RNS). Consistently, we observed increase AMPK phosphorylation at Thr172, along with enhanced phosphorylation of its downstream targets, including ACC, ULK1, and Raptor. This activation leads to the downregulation of mTOR signaling, as evidenced by increased Raptor phosphorylation and reduced phosphorylation of P70S6K, a key mTOR effector. Since mTOR governs anabolic processes such as protein and lipid synthesis and cell growth, its inhibition likely underlies the observed reduction in neuronal soma size. While dysregulation of AMPK and mTOR signaling has been reported in other models of FA (19,43), our study provides direct functional evidence linking AMPK activation to impaired neuronal growth. Supporting this, pharmacological AMPK activation or mTOR inhibition mimicked the soma size reduction in control neurons, while genetic inhibition of AMPK using a dominant-negative AMPKα mutant restored soma size in FXN-deficient neurons. These findings establish AMPK activation as both necessary and sufficient to drive the growth deficit.

Although we cannot fully disentangle the relative contributions of energy depletion and oxidative stress to AMPK activation in this model, both appear to play a role. Beyond sensing ATP levels, AMPK is known to respond to redox signals (44), and ROS/RNS have been shown to activate AMPK and promote compensatory metabolic adaptations such as glycolysis and mitochondrial biogenesis, respectively (45–47). Accordingly, the frataxin-deficient neurons exhibited signs of increased mitochondrial content and glycolytic activity, both hallmarks of AMPK-driven metabolic reprogramming. Moreover, AMPK activation can induce expression of antioxidant response genes, suggesting it may serve as a broader integrator of mitochondrial stress, maintaining cellular homeostasis and promoting cell survival under conditions of stress (48–51).

Together, these findings support a model in which frataxin depletion triggers mitochondrial dysfunction, leading to both ATP depletion and oxidative stress. These stressors converge on AMPK, which in turn inhibits mTOR signaling, thereby impairing neuronal growth. The fact that AMPK activation is reversed by FXN re-expression further reinforces the link between frataxin status and AMPK activation. Ultimately, our data position AMPK-mTOR dysregulation as a key downstream consequence of frataxin deficiency and a mechanistic driver of the soma size phenotype in DRG neurons.

Interestingly, ALA treatment alone decreased AMPK activation, as shown by reduced phosphorylation of AMPKα and its downstream targets. While ALA is traditionally considered a potent scavenger of ROS, our data indicate that its therapeutic action may also arise from restoring mitochondrial function and intracellular ATP levels, thereby indirectly mitigating AMPK hyperactivation. In support of this, ALA was associated with improved mitochondrial energy output, as reflected by reduced mitochondrial oxidative stress and enhanced intracellular ATP levels. Our findings are consistent with previous reports showing that ALA suppresses AMPK activation in human muscles cells (52) and human neuroblastoma cell line (53), likely via indirect effects on metabolic equilibrium rather than through a direct inhibition of the kinase. In our model, we propose that ALA exerts its beneficial effects primarily by enhancing mitochondrial ATP production and reducing ROS, which leads to reduced AMPK activation, normalizes mTOR signaling, and promotes neuronal growth.

Our work establishes a robust and reproducible model of FXN-deficient sensory neurons that captures key cellular consequence of frataxin, including mitochondrial dysfunction, redox imbalance, metabolic signaling alterations, and changes in neuronal morphology. In this contact, we identify a previously unrecognized phenotype of impaired neuronal soma size and link it functionally to dysregulation of the AMPK–mTOR signaling axis. An important strength of this model lies in its scalability and the robustness of its phenotypic readouts. The clear and quantifiable differences between control and KO neurons, combined with its compatibility with multi-well formats (24), make it well suited for phenotype-driven screening approaches. In particular, this system provides a platform to identify compounds that can bypass or substitute frataxin activity or modulate AMPK-mTOR signaling.

However, several limitations should be considered. First, sensory neuron differentiation and subtype representation remain incomplete under current culture conditions. Future refinements, such as co-culture with glia or optimized trophic support may enhance neuronal maturation and better preserve physiological diversity. Second, the use of complete frataxin knockout represents a more severe condition than the partial deficiency observed in FA patients. While this facilitates mechanistic dissection and phenotypic robustness, it may amplify certain cellular response. Complementary models, including human iPSC-derived DRG neuron models carrying GAA expansions (54) will therefore be important to validate the translational relevance.

## Conclusion

In summary, this study defines a cellular framework for investigate the consequence of frataxin deficiency in sensory neurons. We identify AMPK-mTOR signaling as a key mediator linking mitochondrial dysfunction to impaired neuronal growth and establishes ALA as a candidate therapeutic agent capable of restoring bioenergetics and redox balance. These findings contribute to our understanding of proprioceptive neuron vulnerability in FA and pave the way for novel therapeutic approaches that target downstream metabolic pathways in addition to frataxin replacement.

## List of Abbreviations

AAV: Adeno-associated virus
ACC: Acetyl-CoA carboxylase
AICAR: 5-Aminoimidazole-4-carboxamide ribonucleotide
ALA: α-Lipoic acid
AMPK: AMP-activated protein kinase
ATP: Adenosine triphosphate
BCAA: Branched-chain amino acid
CGRP: Calcitonin gene-related peptide
CT: Control
div: Days in vitro
DRG: Dorsal root ganglion/ganglia
Fe-S: Iron–sulfur cluster
FRTL: Ferritin light chain
FXN: Frataxin
GAA: Guanine–adenine–adenine trinucleotide repeat
GFP: Green fluorescent protein
hFXN: Human frataxin
HTS: High-throughput screening
IRP1: Iron regulatory protein 1
KGDH: α-Ketoglutarate dehydrogenase
KO: Knockout
LA: Lipoic acid
LDH: Lactate dehydrogenase
LIAS: Lipoic acid synthase
LplA: Lipoate-protein ligase A
mTOR: Mechanistic target of rapamycin
mtLplA: Mitochondria-targeted lipoate-protein ligase A
NF200: Neurofilament 200
PDH: Pyruvate dehydrogenase
PNs: Proprioceptive neurons
PVALB: Parvalbumin
RPA: Rhodamine-based probe for labile iron
ROS: Reactive oxygen species
SDH: Succinate dehydrogenase
TFR1: Transferrin receptor 1
TMRM: Tetramethylrhodamine methyl ester
ULK1: Unc-51-like kinase 1

## Declarations

## Data availability

All data needed to evaluate the conclusions in the paper are present in the paper and/or the Supplementary Materials.

## Author contributions

Conceptualization, O.G., H.P.; Methodology, O.G., D.M.C., H.P.; Investigation, O.G., D.M.C., A.W., I.M.; Writing O.G., D.M.C., H.P.; Funding Acquisition, O.G., D.M.C., H.P.; Supervision, H.P.

## Acknowledgments

We thank Anne Maglott-Roth, Laurence Reutenauer, Nadège Diedhiou, Aurélie Eisenmann, Valentine Mosbach, and Alain Martelli for technical assistance and discussions. We thank Nadia Messaddeq from the IGBMC imaging platform for EM analysis and Pascale Koebbel (IGBMC molecular biology platform) for vector production.

