## Supplementary data for "Frataxin depletion leads to decreased soma size and activation of AMPK metabolic pathway in dorsal root ganglia sensory neurons"

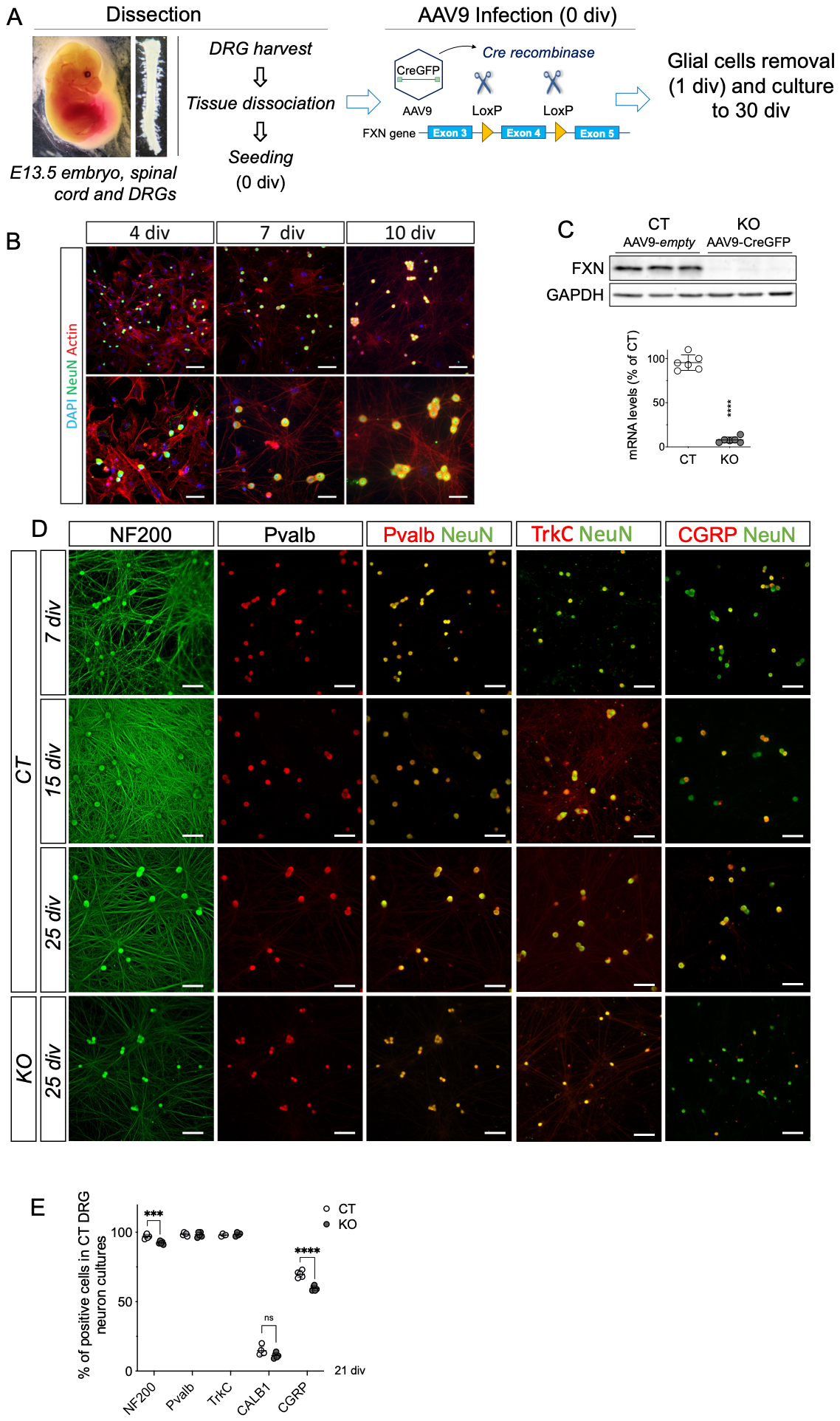

**Figure S1:**

(A) Summary steps of DRG sensory neuron culture model used for these studies. (B) Labeling of nuclei (DAPI), cytoskeleton (filamentous actin) (Phalloidin) and sensory neurons (NeuN) overtime in culture. Glial cells are removed from the culture following antimitotic agent treatment. Scale bars: 100 µm. (C) FXN protein levels (top) and gene expression (bottom) are depleted in AAV9-CreGFP infected cultures (KO), compared to AAV9-empty (control, CT) cultures after 15 div. (D) Sensory neurons fluorescently labeled at 7, 15, and 25 div with antibodies against neurofilament heavy (NF200), parvalbumine (Pvalb), Tropomyosin receptor kinase C (TrkC), calcitonin gene related peptide (CGRP) or pan-neuronal marker NeuN. At Scale bars: 100 µm. (E) Percentage of CT and KO cells positive for NF200, Pvalb, TrkC, CALB1 or CGRP at 21 div. *p<0.05; ****p<0.0001 (Student t-test). Error bars indicate SD.

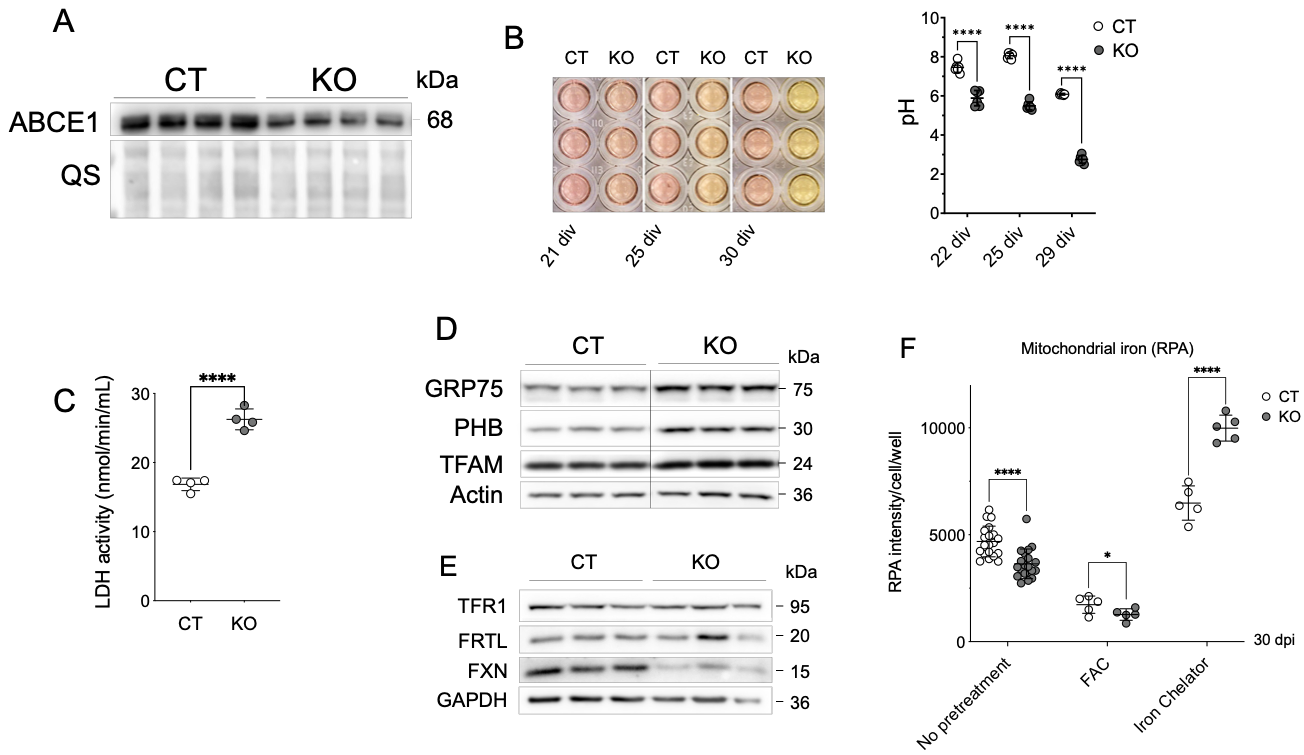

**Figure S2**

(A) Protein levels of ABCE1 (a cytosolic Fe-S protein) assayed by Western Blot in Control (CT) and FXN-depleted (KO) sensory neuron cultures after 21 div. (B) Color and pH quantification of culture media from CT and KO sensory neurons at 21, 25 and 30 div. (C) Lactate Dehydrogenase B (LDHb) activity at 25 div. n = 4. (D) Western Blot of mitochondrial markers Glucose-regulated protein 75 (GRP75), Prohibitin (Phb) and Transcription Factor A Mitochondrial (TFAM) at 30 div. Actin was used as a loading control. (E) Protein levels of iron metabolism related proteins; transferrin receptor 1 (TFR1) and ferritin light chain (FRTL) in CT and KO sensory neurons at 15 div. GAPDH was used as a loading control. (F) Mitochondrial iron content measured at 30 div by RPA sensor in CT or KO cells non pretreated or pretreated for 3 days with 100 µM Ferric Ammonium Citrate (FAC) or 20 minutes with an iron chelator (1,10-phenantroline, 2 mM). RPA signal is inversely correlated to mitochondrial iron levels. Dots indicate averaged RFU (Relative Fluorescence Units) values from every cell of one well (96-well plates, about 300 cells/well). n = 5-10 wells. Error bars indicate SD. (B, C, E) Student t-test. ns = non-significant; *p<0.05; **p<0.01; ***p<0.001; ****p<0.0001. Error bars indicate SD.

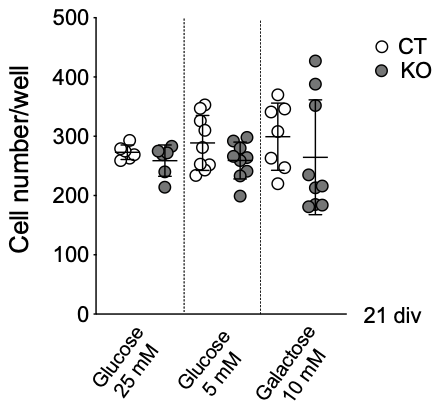

**Figure S3**

Total mitochondrial mass measured at 21 div by Mitotracker Green FM. Dots indicate averaged RFU (relative fluorescence units) values from every cell of one well (96-well plates, about 300 cells/well). Cells were treated with glycolysis inhibitory conditions from 7 div (5 mM glucose, 10 mM galactose). Student t-test. ns = non-significant; **p<0.01; ***p<0.001; ****p<0.0001. Error bars indicate SD.

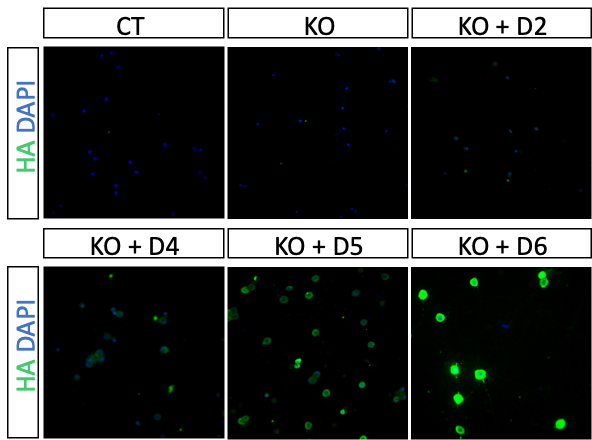

**Figure S4**

Control (CT) and FXN-depleted (KO) sensory neurons were cultured for 7 div, infected with different concentrations of AAV-CAG-hFXN-HA viral particles. (A) Detection of HA-tagged human FXN by immunofluorescence after increasing doses of AAV-CAG-hFXN-HA infection: 3.6 x10^7^ vg/mL (D2) ; 1.8 x10^8^ vg/mL (D4) ; 9 x10^8^ vg/mL (D5) ; 4.5 x10^9^ vg/mL (D6).

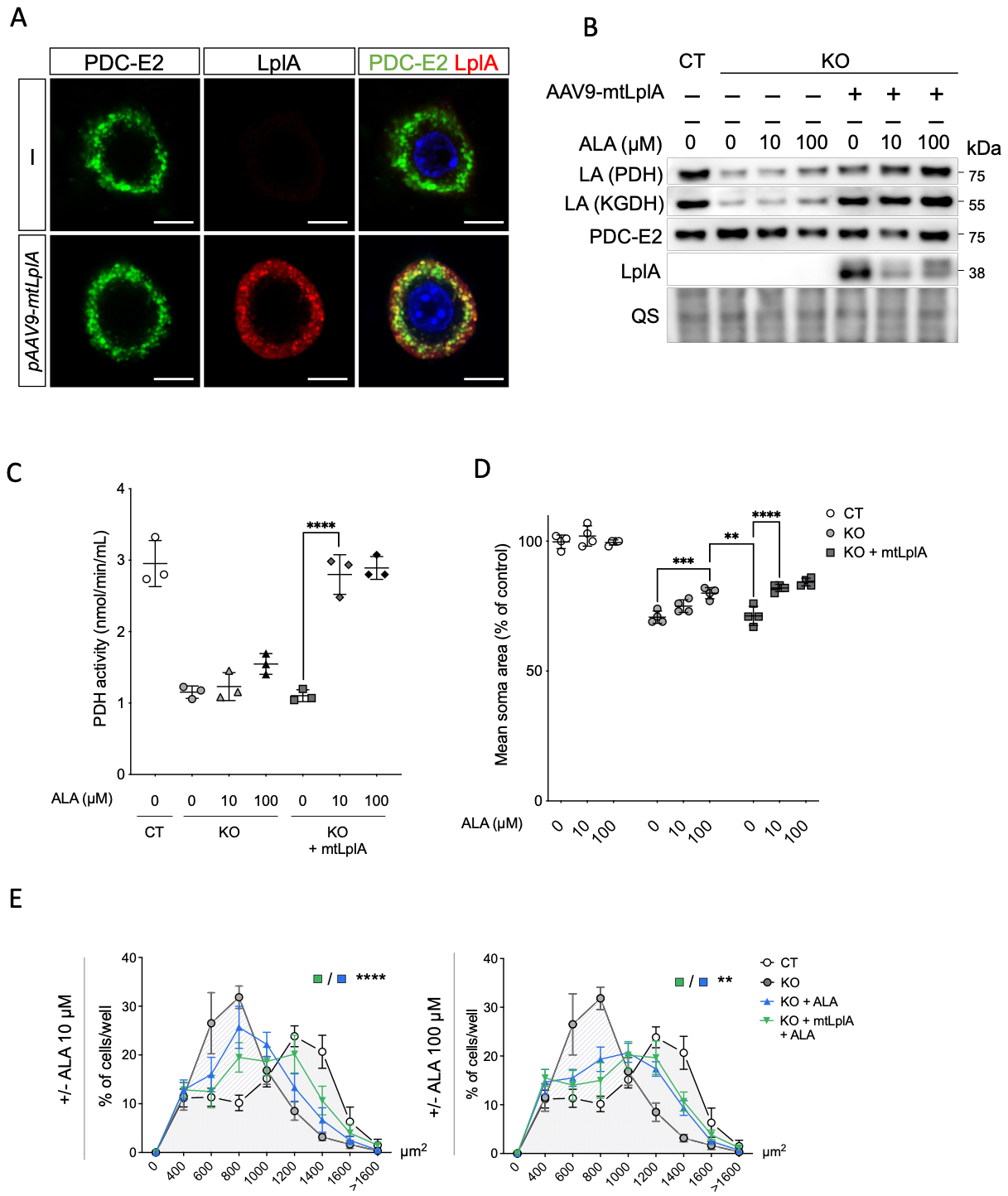

**Figure S5**

(A) Immunofluorescence against LplA (red) and partial co-localisation with mitochondrial marker PDC-E2 (green), in KO neurons infected with AAV9-mtLplA. Cells were further stained with DAPI (blue). Scale bar = 5 µm. (B) Western Blot of Lipoic Acid (LA) on recipient enzymes PDH and KGDH, total PDC-E2, LIAS and FXN at 21 div in CT or KO cells further infected with AAV9-mtLplA and treated by ALA. (C) PDH activity at 21 div of CT and KO neurons infected with AAV9-mtLplA and following alpha-Lipoic Acid (ALA) treatment. (D) DRG neuron soma area in CT, KO or KO + AAV9-mtLplA, measured in 96-well plates when treated with different ALA concentrations. Dots indicate averaged areas from every cell of one well (96-well plates, about 300 cells/well). n = 4 wells. (E) Cells ranked at 25 div by soma areas and frequency of each group calculated (% of cells between N and N-1 size value) in CT and KO neurons infected with AAV9-mtLplA and following alpha-Lipoic Acid (ALA) treatment. n = 6 wells (about 300 cells/well). (B-C) One-way ANOVA followed by Tukey for multiple comparison. (D) Two-way ANOVA with Geisser Greenhouse Correction. *p<0.05; **p<0.01; ***p<0.001; ****p<0.0001. Error bars indicate SD.

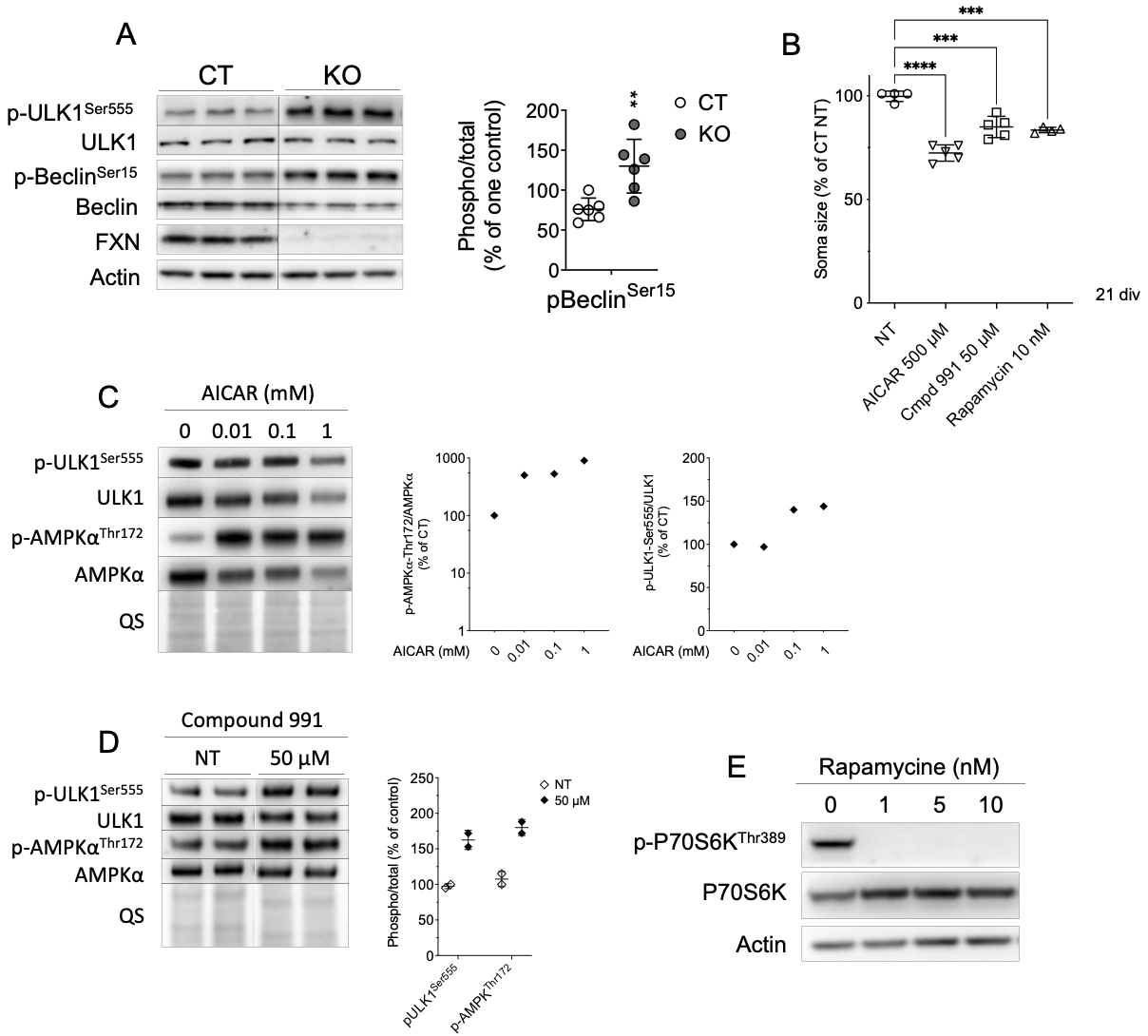

**Figure S6**

(A) Western Blot of ULK1, Beclin and phosphorylated residues at Serine 555 and Serine 15, respectively. Right panel shows quantification of phospho-Beclin^Ser15^ relative to total Beclin. Actin was used as a loading control for quantifications. Student t-test. **p<0.01. Error bars indicate SD. (B) Measurement of soma size in 21 div CT cultures untreated (NT) or treated for 14 days with 500 µM AICAR, 50 µM Compound 991 or 10 nM Rapamycin. n = 5-10 wells (about 300 cells/well). One-way ANOVA followed by Tukey for multiple comparison; ****p<0.0001. Error bars indicate SD. (C, D) Western Blot of phosphorylated and total proteins of AMPK pathway at 21 div following 96h treatment with different concentrations of AICAR (C) or 50 µM Compound 991 (D). Right panels indicate quantification of the Western Blot. QS staining was used as a loading control for quantifications. (E) Western Blot of phosphorylated and total proteins of mTOR pathway, following 96h treatment with different concentrations of rapamycin. Actin was used as a loading control.

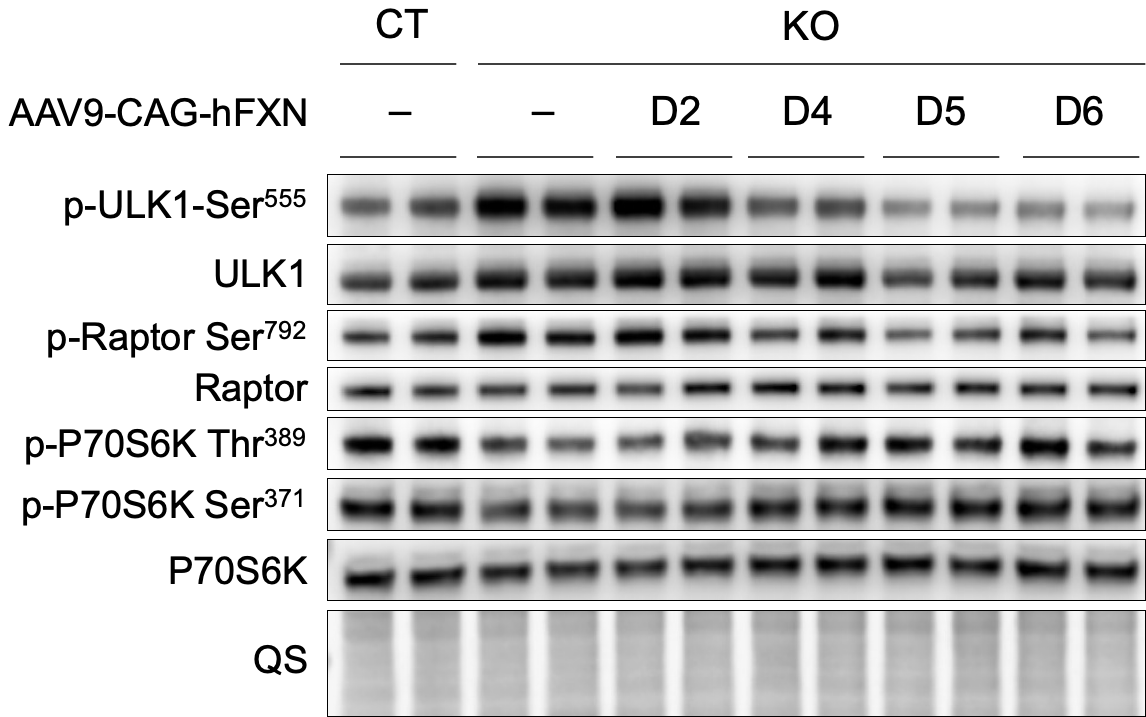

**Figure S7**

Western blot analysis of phosphorylated and total proteins of AMPK and mTOR downstream targets, after increasing doses of AAV-CAG-hFXN-HA infection: 3.6 x10^7^ vg/mL (D2) ; 1.8 x10^8^ vg/mL (D4) ; 9 x10^8^ vg/mL (D5) ; 4.5 x10^9^ vg/mL (D6).

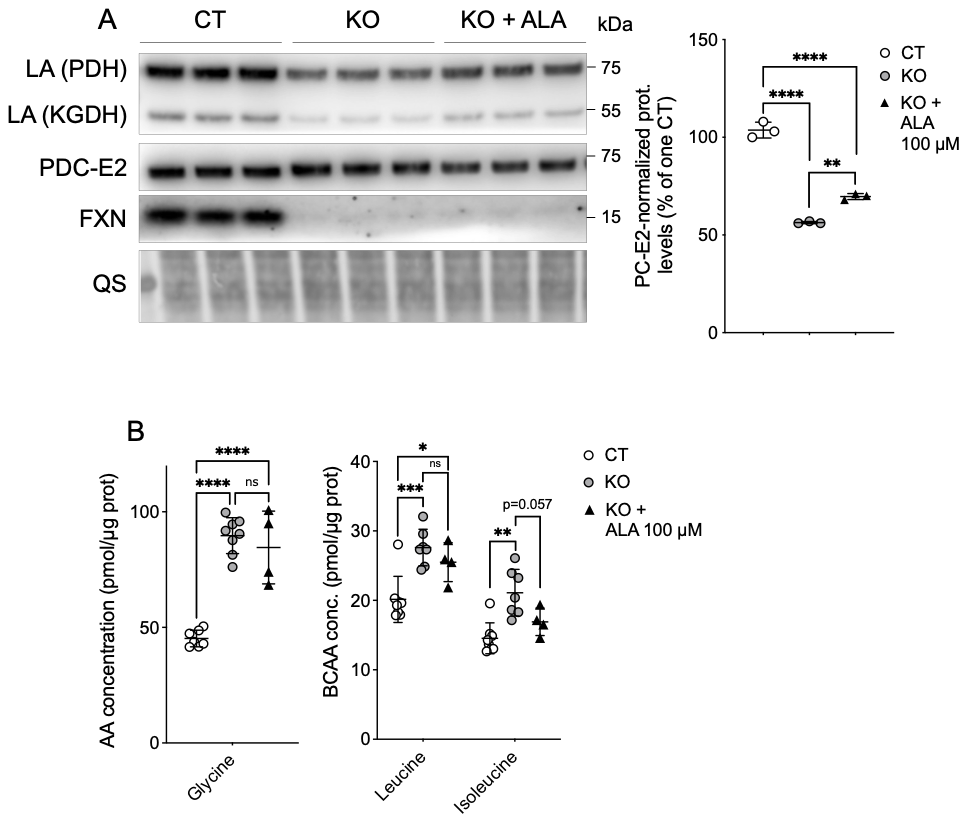

**Figure S8**

(A) Western Blot of Lipoic Acid (LA) on recipient enzymes PDH and KGDH, total PDC-E2 and FXN at 21 div. QS staining used as a loading control for quantification. (B) Measure of concentration of glycine, leucine and isoleucine in CT and KO cells at 30 div, with or without ALA treatment. One-way ANOVA followed by Tukey for multiple comparison. *p<0.05; **p<0.01; ***p<0.001; ****p<0.0001. Error bars indicate SD.

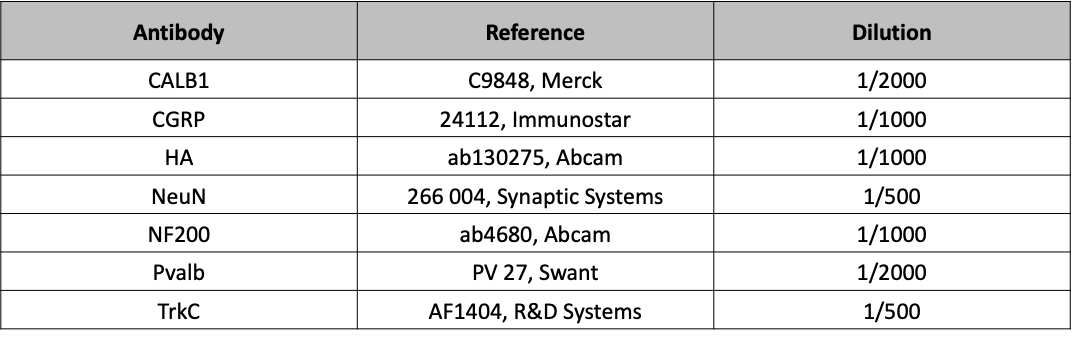

**Table S1**

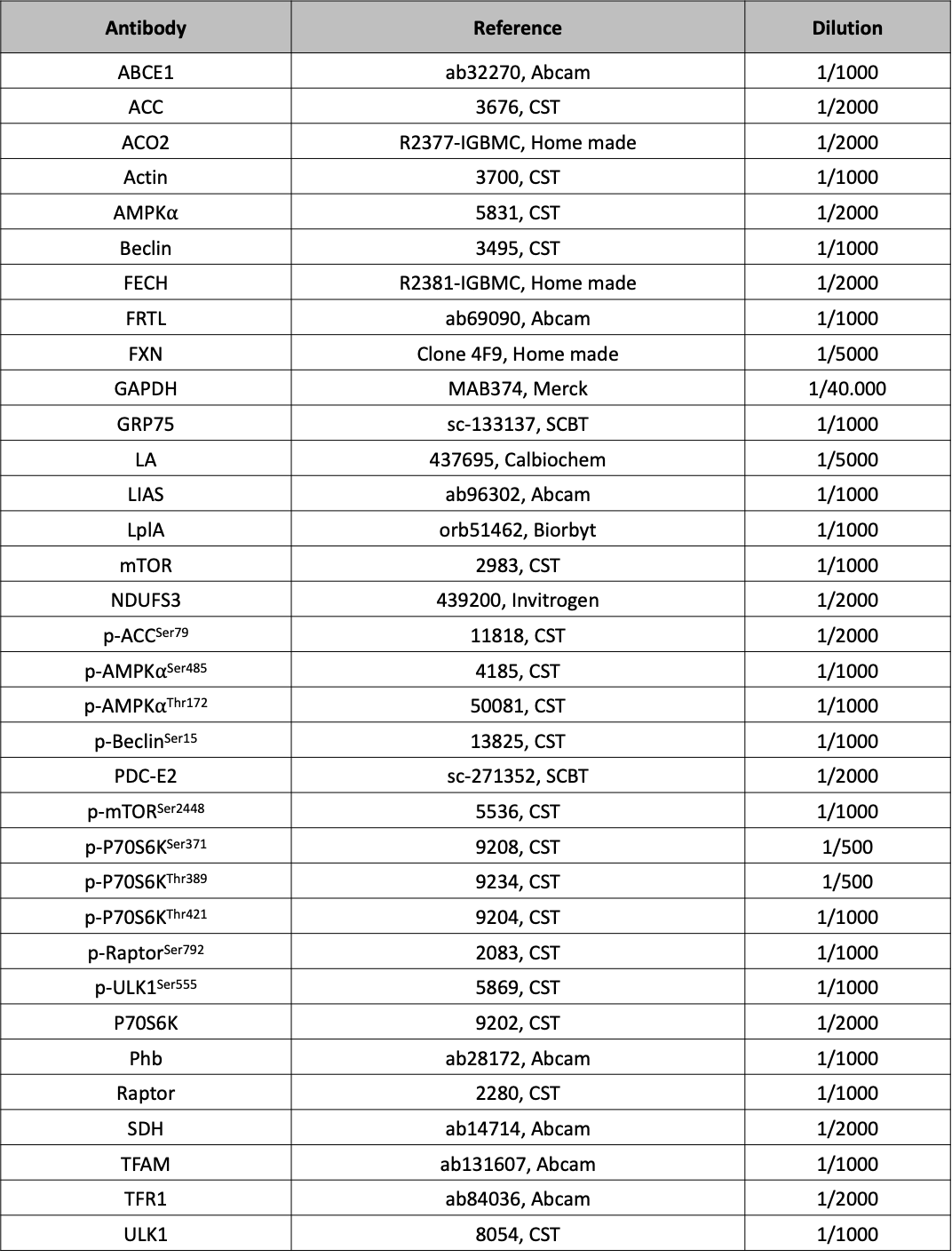

**Table S2**

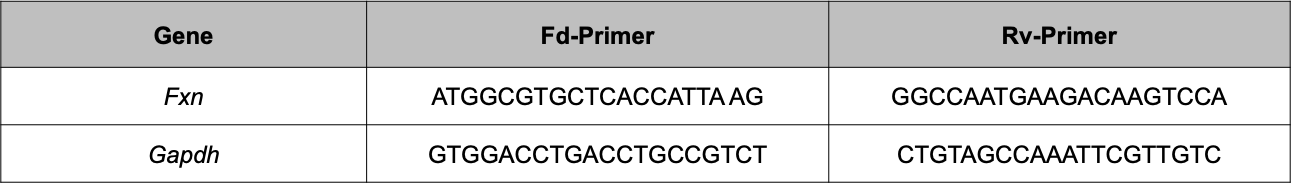

**Table S3**
